# Elucidating Electronic Structure Variations in Nucleic Acid-Protein Complexes Involved in Transcription Regulation Using a Tight-Binding Approach

**DOI:** 10.1101/2024.04.15.589549

**Authors:** Likai Du, Chengbu Liu

## Abstract

Transcription factor (TF) are proteins that regulates the transcription of genetic information from DNA to messenger RNA by binding to a specific DNA sequence. Nucleic acid-protein interactions are crucial in regulating transcription in biological systems. This work presents a quick and convenient method for constructing tight-binding models and offers physical insights into the electronic structure properties of transcription factor complexes and DNA motifs. The tight binding Hamiltonian parameters are generated using the random forest regression algorithm, which reproduces the given *ab-initio* level calculations with reasonable accuracy. We present a library of residue-level parameters derived from extensive electronic structure calculations over various possible combinations of nucleobases and amino acid side chains from high-quality DNA-protein complex structures. As an example, our approach can reasonably generate the subtle electronic structure details for the orthologous transcription factors human AP-1 and Epstein-Barr virus Zta within a few seconds on a laptop. This method potentially enhances our understanding of the electronic structure variations of gene-protein interaction complexes, even those involving dozens of proteins and genes. We hope this study offers a powerful tool for analyzing transcription regulation mechanisms at an electronic structural level.

**Topic of Content:** Transcription factors that bind to DNA modulate gene expression, with the stability and reactivity of their interactions elucidated by eigenvalues derived from the tight-binding model. Visualization of these interactions reveals the Highest Occupied Molecular Orbital (HOMO) and the Lowest Unoccupied Molecular Orbital (LUMO), the gap between which determines the reactivity and stability of the molecular complex. This approach advances our understanding of gene regulation by revealing the dynamics of charge transfer and electronic states within transcription factor-DNA complexes.

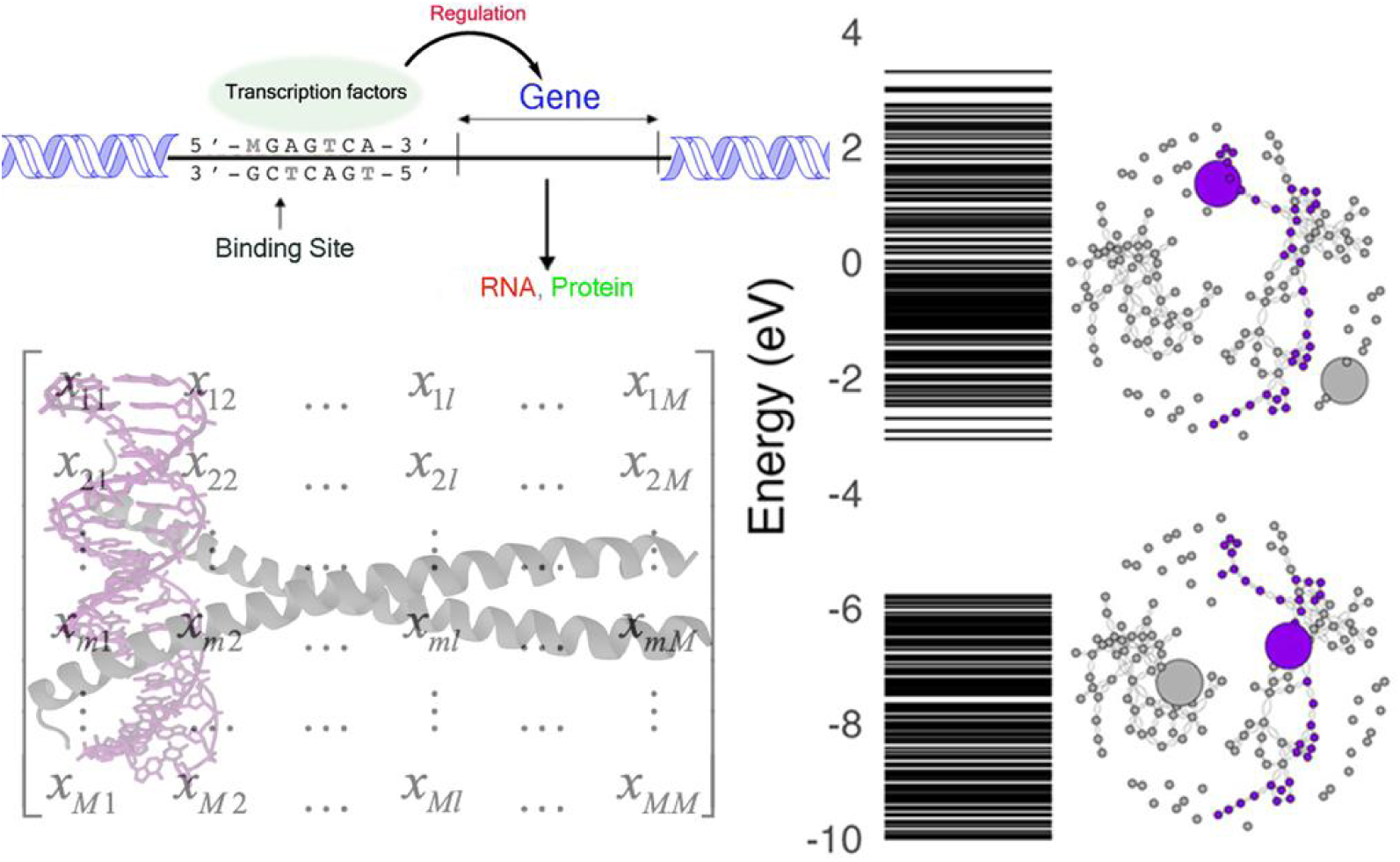

## Introduction

Protein-DNA interactions play a crucial role in various biological processes, such as gene regulation, transcription, DNA replication, repair, and packaging.(1–4) For decades, the quest to understand the intricate relationships between DNA and proteins has been at the heart of biological research.(5–10) These nucleic acid-protein interactions usually occur in two ways: non-specifically, such as the interaction between histones and DNA, and through highly selective, sequence-specific binding, as seen in transcription factors. This distinction is essential for numerous biological functions, ranging from gene regulation to DNA repair.(11) Eukaryotic DNA is packaged into nucleosomes (Figure 1).(12–15) The nucleosome core particle (NCP) is the fundamental unit of DNA packing in eukaryotic cells. It consists of an octamer of histone proteins around which approximately 150 base pairs of DNA are bound.(16–18) The fundamental unit of DNA packing inside eukaryotic cells is the nucleosome core particle (NCP), in which approximately 150 base pairs of DNA are bound around an octamer of histone proteins. Transcription factors (TFs) act as mediators of genetic information, directing the complex process of transcription, in which DNA is transcribed into RNA, a precursor to protein synthesis.(6–10, 19–21)

**Figure 1.**
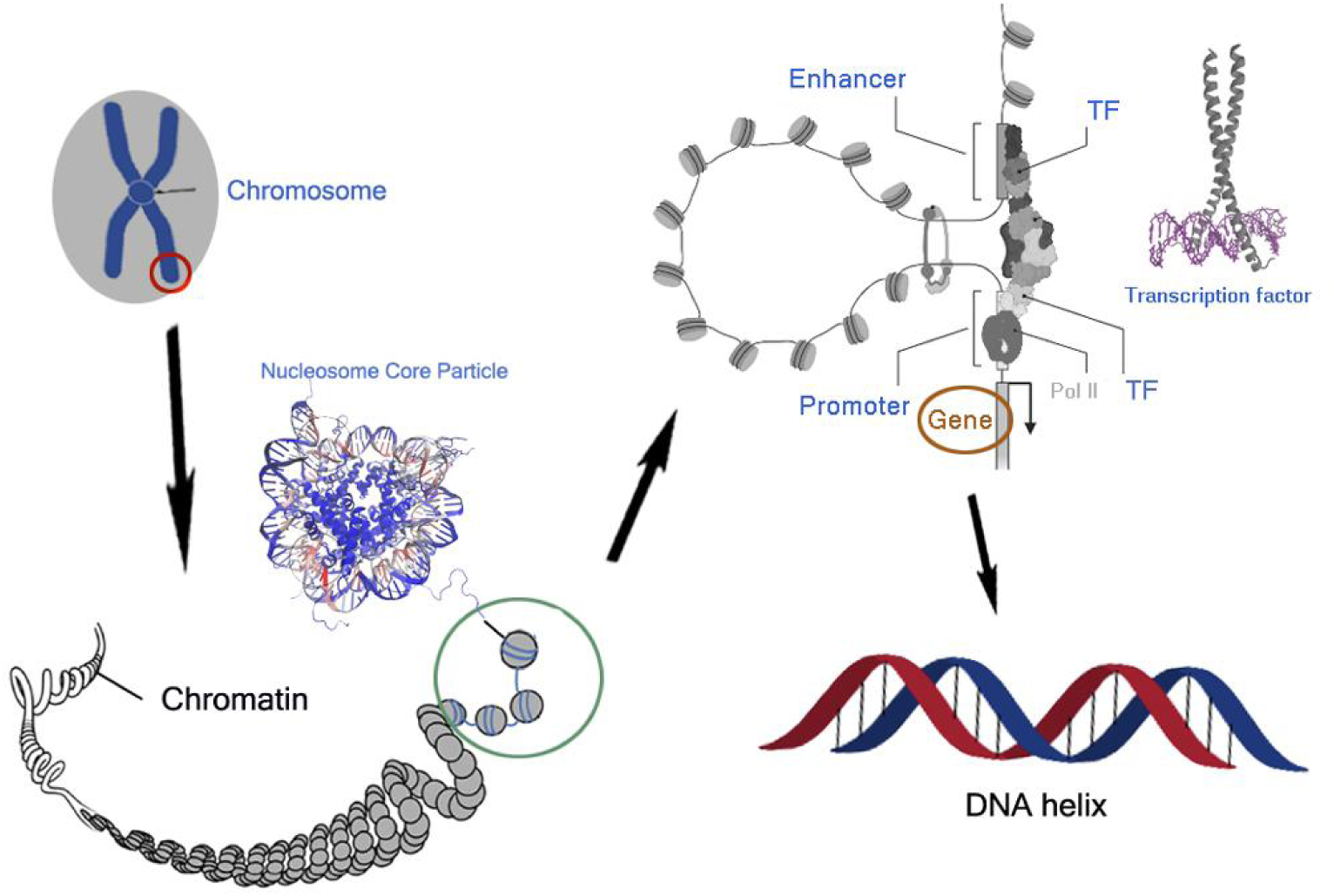
The hierarchical structure of the chromosome organization with emphasis on transcriptional regulation, starting from the chromosome level, through chromatin and the nucleosome core particle, to the DNA helix. The atomic-resolution structures of the NCP and TF are also given.

The activator protein-1 (AP-1) is a regulatory element that is present in many promoter and enhancer regions. AP-1 plays a crucial role in regulating gene transcription across various biological functions, highlighting its versatility in cellular biology.(22–25) And it is characterized by the presence of a highly conserved DNA binding domain that contains an N-× 7-R/K sequence and a basic leucine zipper (bZip) domain.(26–32) The relatively poorly conserved leucine zipper region is characterized by leucine in the last position of every seven amino acids, and hydrophobic residues.(28, 33, 34) AP-1 proteins are a versatile family of dimeric transcription factors. Jun protein is a member of the AP-1 proteins. It has the ability to form homodimers or heterodimers with other proteins. The c-Jun protein promotes cell cycle progression by repressing the p53 tumor suppressor and activating cyclin D1. This reduces the influence of the cyclin-dependent kinase inhibitor (CDKI) p21, facilitating the G1 to S phase transition.(35–38)

Exploring the impact of electron injection on DNA-binding proteins is important in various research fields. Ultrafast electron transfer occurs during the recognition of various DNA sequences by a DNA-binding protein with distinct dynamic conformations.(39–44) DNA damage and repair mechanisms involve electron transport. For instance, positive charge transfer can promote oxidative damage to guanine in DNA, which may be related to the presence of mutation sites in the genome.(45–54) DNA transcription factors such as SoxR and p53, which are equipped with redox-active groups, use DNA charge transport as a redox sensing mechanism.(55–58) The DNA-mediated charge transport might enable signaling between the [4Fe4S] clusters in the human DNA primase, polymerase α, and other replication and repair high-potential [4Fe4S] proteins.(59–63) This DNA charge chemistry serves as both a sensing method and a monitor of DNA integrity, which is sensitive to base stacking perturbations caused by mismatches or DNA damage.

Quantum chemistry provides chemists with critical insight into the electronic structure behavior of DNA or protein molecules, but its extensive computational requirements limit the scope and variety of systems that can be effectively analyzed.(64–68) The tight-binding (TB) method offers a more practical alternative for describing the electronic Hamiltonian using smaller and more sparse matrices.(69–74) In early work, the TB model was applied to materials science or solid state physics. The TB model has been applied to molecular clusters or biomolecular systems.(75–81) Traditionally, the TB Hamiltonians have relied on empirical or semi-empirical parameters, which raises concerns about their accuracy and general applicability.(82–89) A few works are developed to improve the accuracy and dependability of TB models through the foundation of first-principles calculations.(90–92)

The Protein Data Bank (PDB) has provided a continuous influx of high-resolution structural data, which has significantly advanced our understanding of protein-DNA interactions.(93–96) The increasing number of high-quality experimental protein and DNA structures, including those obtained through X-ray, NMR, and cryo-EM techniques, have provided opportunities to improve our TB parameters for biological systems. As previously proposed, it is possible to derive TB parameters for millions or even billions of molecular fragments, which represent most occurrences in protein and DNA databases(92, 97). Integrating structural insights, especially regarding residue preferences in protein-DNA interactions, is essential for understanding charge transfer mechanisms. Although accuracy is improved, constructing the Hamiltonian is time-consuming due to the cost of ab initio calculations and the projection step. Furthermore, the resulting *ab initio* TB Hamiltonian is not transferable to new structural configurations, which limits its usefulness for electronic structure simulations. Nowadays, machine learning algorithm in computational chemistry(98–106) has been widely used to predict interaction energies, molecular forces(107, 108), electron densities(109), density functionals(110) and various molecular response properties(111–114) The machine learning algorithm can be used to predict accurate TB Hamiltonian for unseen structures during atomic structure explorations. Therefore, the machine learning method for TB Hamiltonian parameterization is desired.

In this work, we investigate DNA-protein interactions in transcriptional regulation with a focus on transcription factors, which regulates the transcription of genetic information from DNA to messenger RNA by binding to a specific DNA sequence. A comprehensive library of residue-level tight binding parameters is constructed from detailed electronic structure calculations. The library covers millions of nucleic base and amino acid side-chain combinations extracted from high-quality DNA-protein complex structures. TB Hamiltonian parameters derived from *ab-initio* calculations are accurately generated using a random forest regression algorithm. Despite its simplifications, the direct diagonalization of the TB Hamiltonian could generate various electronic structure properties of DNA-protein complexes. Our approach quickly reproduces the electronic structure details of orthologous transcription factors, such as human AP-1 and Epstein-Barr virus Zta(115, 116), in seconds using a laptop. We anticipate that our study will serve as a powerful tool for analyzing transcription regulation mechanisms at an electronic structural level. And this methodology opens up possibilities for comprehending the electronic structure variations observed in millions of protein-gene complexes or dozens of gene-protein complexes, in the big data scenario.

## **2.** Methods and Computational Details

### Construction of the Nucleobase-Amino Acid Library

The DNA-protein complexes contain only the twenty L-amino acids and four deoxynucleotides, which are generally distinguished by their different side chain structures and chemical compositions (Figure 2). DNA-backbone interactions are the most numerous and contribute to the stability of the DNA-protein complex. In contrast, side-chain interactions of the protein are fewer but confer specificity by recognizing the unique features of the DNA sequence. The TB parameter library currently includes collections of all possible combinations of amino acids and nucleobases, specifically the amino acid/amino acid (AA), base/base (BB), and amino acid/base (AB) interaction patterns. Our previous work(92, 97, 117) has thoroughly studied the AA and BB conformers, so this study will focus solely on the AB conformers. Note that the BB conformers in previous work were generated from customized DNA models using packages such as x3DNA(94). In this work, we have updated the BB conformers based on experimental DNA protein structures. The procedure to extract each conformer from the available three dimensional DNA binding protein structures follows the work of Singh and Thornton(118). This library comprises around 1.2 million conformers that cover a broad range of nucleic acid sequences and protein families, ensuring representation across different binding modes. The initial structures in the library only contain the coordinates of the heavy atoms. The missing hydrogen atoms were added using the *tleap* module in the AmberTools package(119). Three protonation states were calculated for histidine, and two possible protonation states were considered for other acidic and basic amino acids.

**Figure 2:**
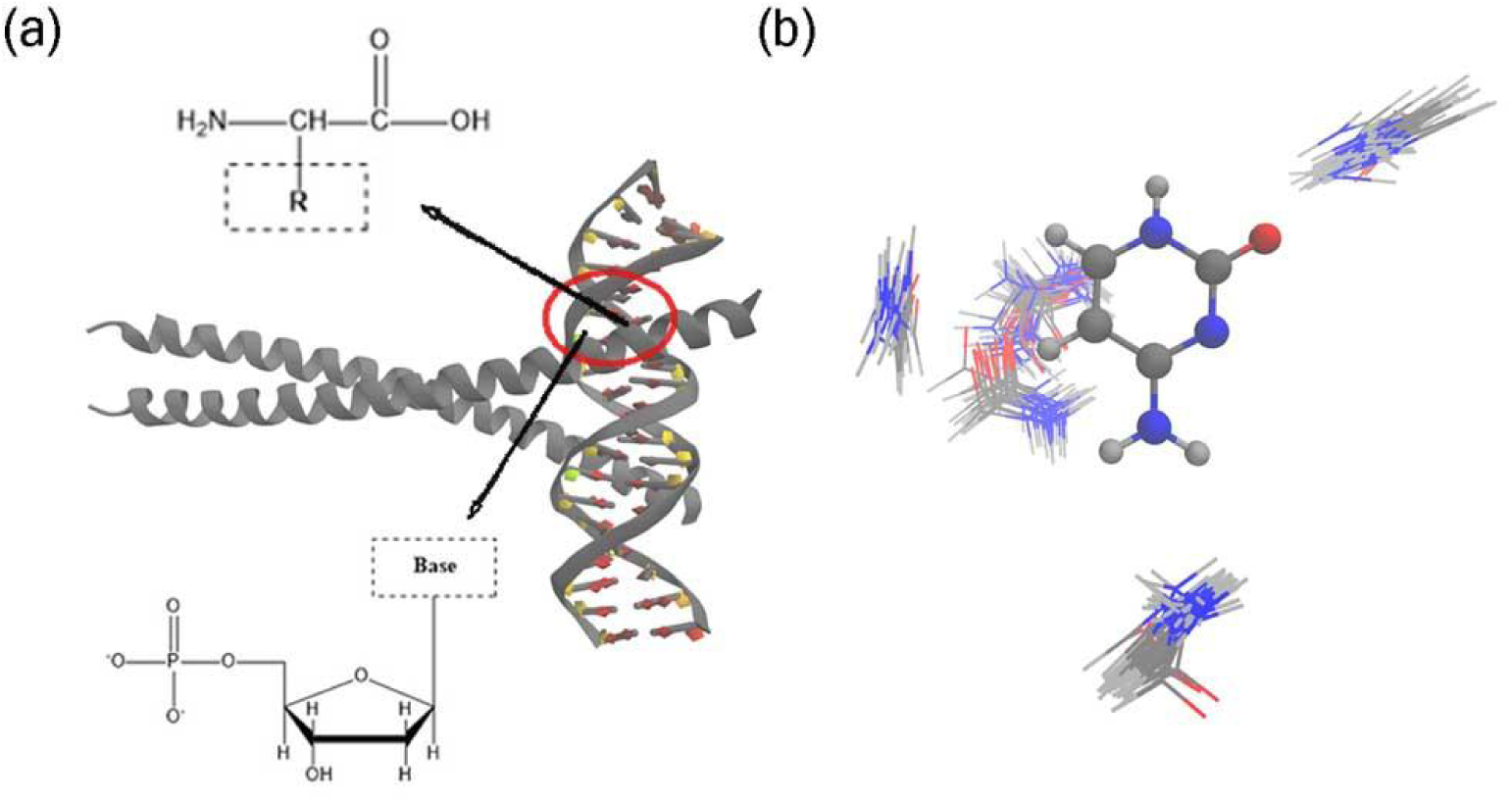
Illustration of one of the studied nucleobase-amino acid system (PDB ID: 2H7H). (b) Depiction of the nucleobase’s phosphate group linked to a sugar ring, which in turn is bonded to a base. Adjacent is the general structure of an amino acid, with its variable side chain represented by "R" in a dashed outline. (b) Spatial distribution patterns of the interactions between the cytosine base (CYT) from the nucleotide and the glutamine (GLU) side chain. The clusters highlight various conformers.

### The Data Driven Tight Binding Model for Biomolecules

The tight binding model is a robust framework for studying the electronic properties of large and intricate molecular systems. The foundational principles of the tight binding model for molecule systems, including the derivation process, have been detailed in previous publications from us(92, 97, 117, 120) or contributions by others(121, 122). Here, we only describe our methodologies for calculating on-site energies, charge transfer couplings, and the Löwdin transformation in our current research.

Biomolecules are composed of repeated structural units, such as amino acids for proteins and nucleotides for DNA. In the tight-binding approximation, electrons have limited interactions with non-neighboring sites. The formulas for on-site energy and transfer integral are provided below:

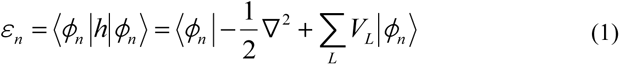

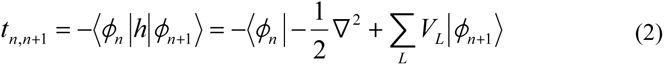

The summation runs over all possible sites *L*. However, only the neighboring sites need to be considered in the TB approximation. And *ε* represents the on-site energy and *t* represents the transfer integral between sites. *ϕn* refers to the molecular orbital of one structural unit *n*. Therefore, the on-site energy for site *n* only requires the potential information of site n and its closest neighboring sites ***C***. The formula for on-site energy can be simplified as follows:

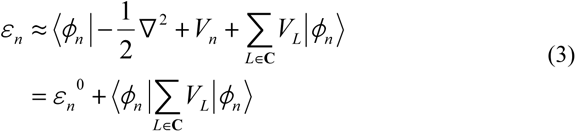

According to Equation 3, the on-site energy is not solely determined by the orbital energy of site *n*; it also includes contributions from adjacent sites, particularly the first set of nearest neighbors, denoted as ***C***. The model can take into account the impact of neighboring residues on the on-site energy.

The transfer integral describes the ability to perform charge transfer among neighboring sites, while the on-site energy describes the ability to move or inject an electron from a specific site. The transfer integral only require the potential of site *n* and *n*+1, that is

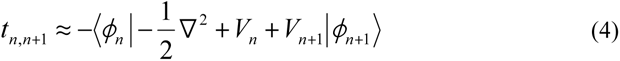

In this work, we utilize the Löwdin method to minimize orbital overlap, as the tight binding model corresponds to the orthogonal basis. This enables us to transform the effective transfer integral.

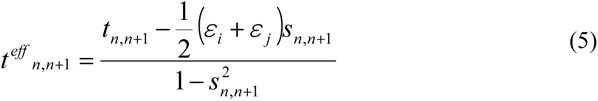

Equation 5 defines *s* as the orbital overlap integral between sites. This transformation has minor effects on the on-site energy and can be safely ignored if necessary. The TB parameters have been extensively studied for pure DNA complexes and protein complexes in the previous work(92, 123).

In the framework of the tight binding Hamiltonian, the on-site energy and transfer integrals are characterized as the diagonal and off-diagonal matrix elements, respectively. Diagonal elements correspond to the on-site energy for a given orbital or site, which signifies the energy level of an electron when it is localized at that site. Conversely, off-diagonal elements quantify the transfer integral, indicative of the probability of an electron’s transition between sites, which is a measure of the charge transfer couplings within the molecular system.

Another practical difficulty is the inefficiency in constructing the TB Hamiltonian from *ab initio* calculations. Here, the random forest (RF) regression is utilized to predict TB parameters within the BioTinter-1m framework. The RF regression model is employed as a multi-input and multi-output framework(124–129), enabling the simultaneous prediction of all TB parameters. This method constructs an ensemble of decision trees from varied segments of the training data, enhancing model diversity and robustness. Each decision tree’s construction is guided by random subsets of features, enabling nuanced learning from the dataset. The RF model averages predictions across all trees to estimate molecular descriptors, as implemented in the scikit-learn module(130) in Python. The ensemble of 150 trees balances computational efficiency with predictive accuracy.

Although various machine learning techniques were explored, including deep learning methods(131–136), the findings indicate that the performance of deep neural networks does not surpass that of the RF model. The limited success observed in our studies with deep neural networks can often be attributed to insufficient data in the training set. Although our library contains millions of biomolecular residues, only a few hundred or thousand conformers are available for each type of AA, BB, or AB combination. Our initial test with the deep neural network model implemented in PyTorch resulted in a correlation coefficient below 0.92 and was therefore not reported. In contrast, the RF model showed the lowest correlations of 0.95 or higher (see Table S1). Expanding the dataset by a factor of 100 or 1000 could potentially enhance the predictive capability of deep learning networks and improve the overall understanding of biomolecular electronic structure variations. In our preliminary evaluations, the deep neural network model, implemented using the PyTorch framework(137), exhibited the correlation coefficient of less than 0.91, which did not meet our benchmark criteria for inclusion in this study. The RF model demonstrated relatively superior performance, consistently achieving correlation coefficients of 0.95 or above, as detailed in Table S1. We hypothesize that augmenting our dataset by an order of magnitude, specifically by factors of 100 to 1000, might significantly enhance the ability of deep neural network to predict and thereby offer more profound insights into the variability of electronic structures in biomolecular systems.

After constructing the TB Hamiltonian, we can solve the well-known eigenvalue equation (**HC**=**EC**) directly for electronic structure calculations of any bio-molecules. The electron-ion dynamics can also be solved within the TB framework. These methods are implemented in our in-house code BioTinter (**T**ight-binding model for **Bio**molecular ***inter***actions). Because this code carries a TB parameters library of 1.2 million conformers, we would also refer to it as BioTinter-1m. The workflow of BioTinter is shown in Scheme 1.

**Scheme 1.**
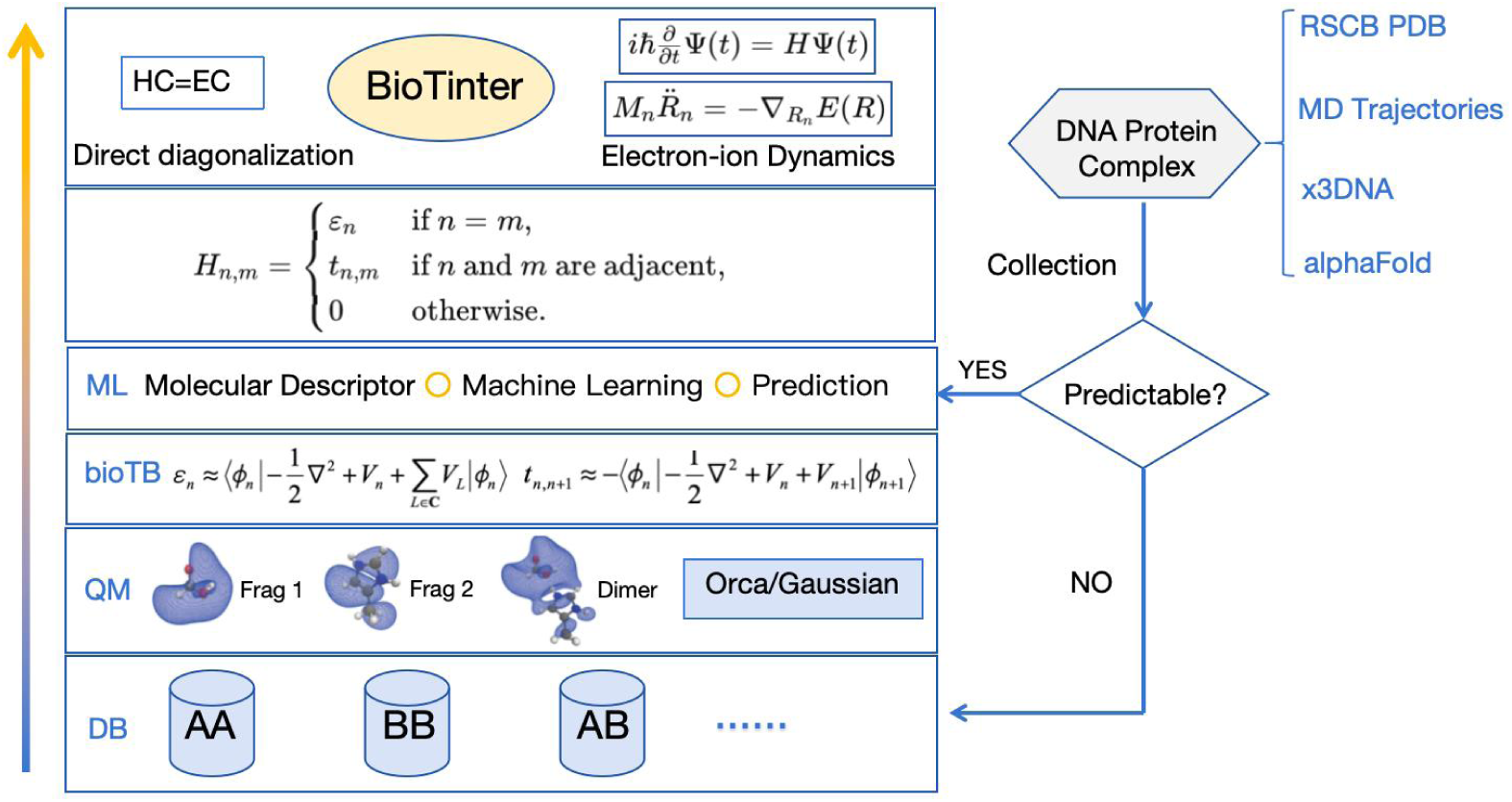
The workflow and code structure of BioTinter package used in this work.

The BioTinter framework employs a layered architecture to integrate TB parameters into quantum chemistry workflows, significantly enhancing the computational efficiency and accuracy of molecular simulations involving DNA-protein complexes. At its core, the Database (DB) layer hosts an extensive library of pre-calculated TB parameters. Absent parameters trigger the Quantum Mechanics (QM) layer, which calculates needed parameters via interfaces with Orca(138) and Gaussian(139) to compute the requisite parameters. This process is augmented by the bioTB module, as detailed in our preceding publications(92, 97). The Machine Learning (ML) layer predicts TB parameters for novel conformers, enabling the construction of the TB Hamiltonian for simulations. Initial structural data for simulations are sourced from the Protein Data Bank (PDB), MD trajectories, or tools like x3DNA(94) and AlphaFold(140). BioTinter-1m prioritizes a balance between speed and accuracy, resorting to on-the-fly QM calculations when necessary. This on-the-fly module ensures that even with a vast database, the system remains responsive and accurate. The upcoming public release of BioTinter-10b may weaken this on-the-fly module, as the conformer library is expected to expand to ten billion entries along with deep neutral network model.

### Simulation Details

In order to construct the TB parameters library, the positions of hydrogen atoms were optimized for each dimer using B3LYP/6-31G(d) calculations. We kept the coordinates of the heavy atoms fixed during the optimization process. The on-site energies and charge transfer couplings for each dimer are derived from at the HF/6-31G(d) and B3LYP/6-31G(d) level according to the idea of tight-binding approximation as our previous work.(92, 97) The solvent effects were considered with the implicit solvation model if necessary. Quantum chemistry calculations can be performed using either the Gaussian or Orca package, both of which have been interfaced with BioTinter.

In the ML layer, the relative positions of molecules are described through their internal coordinates (IC), the Coulomb Matrix (CM) and Smooth Overlap of Atomic Positions (SOAP) descriptors. For a comprehensive understanding of CM and SOAP descriptors, we recommend referring to existing literature.(141–144) Our analysis considers the effect of including or excluding hydrogen atoms in these molecular representations. Benchmark results (Table S1) reveals that presence of hydrogen atoms does not significantly affect our model’s predictions. This research primarily uses hydrogen-depleted CM descriptors, which are refined using a norm sorting technique. While the SOAP model introduces a more complex approach, it only slightly improves predictive accuracy. Therefore, our approach in BioTinter-1m prioritizes hydrogen-depleted CM descriptors for simulating DNA-protein systems.

To illustrate the utility of the tight-binding (TB) model, we investigate the electronic structure variations in complexes involving Activator Protein 1 (AP-1) and Epstein-Barr Virus Zta transcription factors with their associated nucleic acids. The coordination of this sophisticated computational process is facilitated by the Snakemake workflow management system(145, 146). Calculations are monitored and streamlined using custom Python scripts developed for the BioTinter packages, ensuring an automated and efficient workflow. Subsequent statistical analysis of the results is performed using R scripts, providing a comprehensive assessment of the models’ predictive accuracy.

## Results and Discussions

TB parameters were calculated for thousands of AB conformers to analyze the specialization of amino acid or nucleic base distributions in realistic DNA-protein complexes. A complete tight binding Hamiltonian can be constructed for any DNA-protein complex by combining previously reported TB parameters from AA and BB libraries(92, 97). After collecting the AA, BB, and AB distributions, there are approximately one million conformers. This library is useful for describing how the conformation ensemble influences TB parameters within distinct protein structures. For instance, the TB parameters library allows for the extraction of explicit geometric correlation with the charge transfer couplings. It is commonly observed that the values of the charge transfer couplings rapidly decay, decreasing to negligible levels at distances closer than 6.0 Å.

The principal component analysis (PCA) algorithm was used to categorize various AB parameters and correlate them with their physical properties. Figure 3 displays a two-dimensional (2D) plot from PCA that separates the data into distinct clusters. The color coding represents different amino acid characteristics: acidic (red), basic (blue), hydrophobic (purple), and polar (gray), highlighting the chemical nature of the residues as a pivotal factor in the variability of tight binding parameters. The numbers in the brackets on the PC1 and PC2 axes of the PCA plot represent the percentage of the variance in the dataset that is explained by each principal component. This plot also demonstrates the intrinsic distribution of parameters within each cluster, distinctly influenced by nucleobase type—adenine (ADE), thymine (THY), guanine (GUA), and cytosine (CYT). To ensure functional selection independence, TB parameters were calculated using the Hartree-Fock (HF) method. For comparison, TB parameters were also calculated using the B3LYP level method, as shown in Figure S1. The PCA plots resulting from the B3LYP calculations confirm the segregation of data into distinct clusters, as observed with HF calculations. The spatial arrangement of TB parameters in AB conformers is primarily determined by the chemical nature and charge state of the amino acid residues. Secondary factors include the type of nucleobase and the choice of DFT functional.

**Figure 3.**
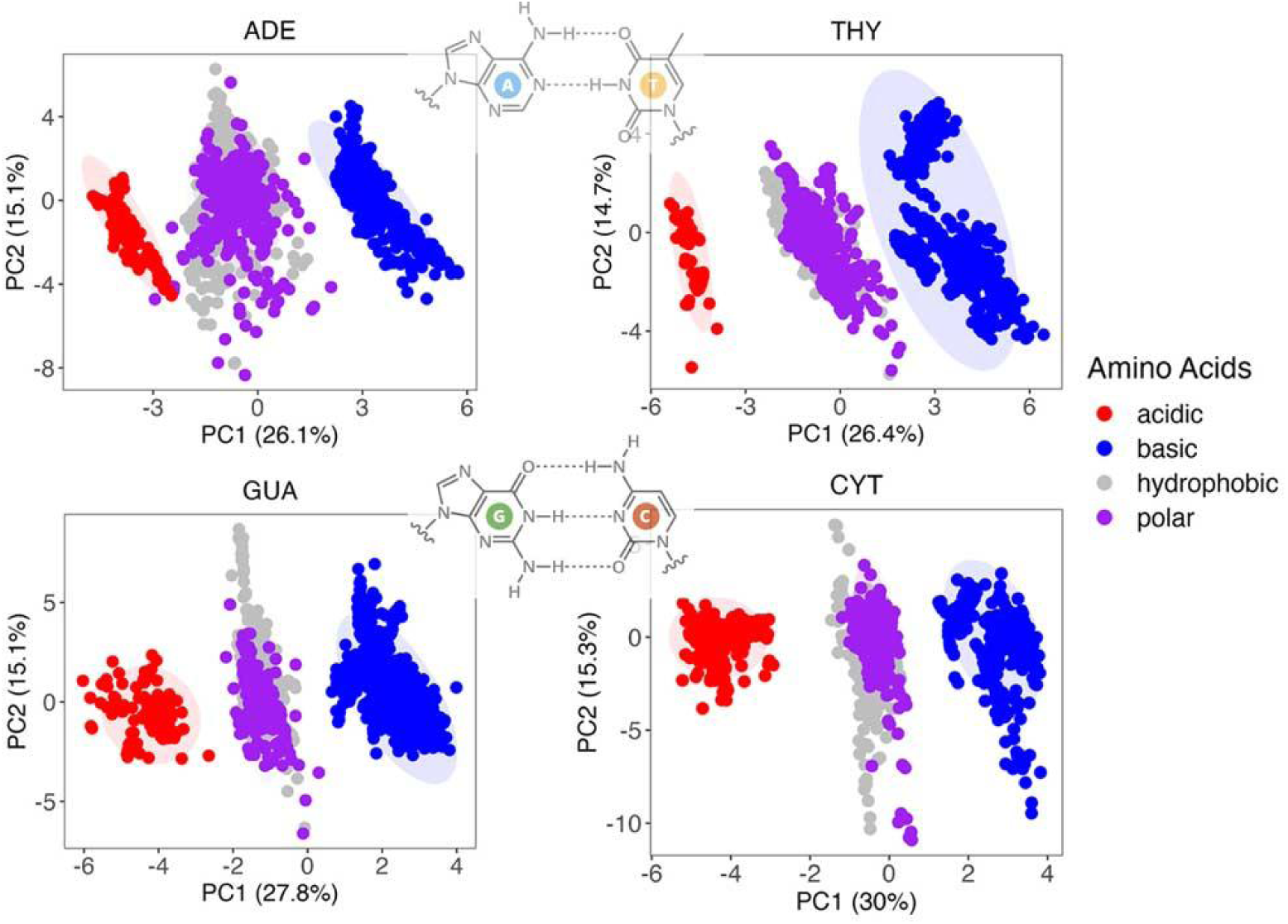
The PCA visualization of a spectrum of TB parameters involving HOMO and LUMO orbitals. The visualization includes the four types of nucleic bases, which are the components for any possible DNA sequence. The confidence ellipse represents a statistical probability of 95% that encloses a certain percentage of the data points based on their distribution along the principal components.

Figure 4 shows a detailed analysis of the average hopping integrals between each of the four nucleobases and twenty standard amino acids. This figure also highlights the varying interaction strengths of histidine in its three protonation states: HID, HIE, and HIP, which reflect the different coupling strengths in various biochemical environments. The charge transfer integrals between nucleobases and various amino acids exhibit significant differences. Each nucleobase has its own preferred interacting amino acid with specific charge transfer couplings. This is fundamental in comprehending the dynamics of DNA-protein interactions at the electronic and molecular levels. Aromatic amino acids, such as histidine, phenylalanine, tryptophan, and tyrosine, generally exhibit significant charge transfer couplings. This phenomenon may be caused by either the π-π interaction or the C-H-π interaction, which could significantly enhance the possibility of electron transfer. The average on-site energy difference for such AB conformers is often within 1.0 eV or even lower. Other residues, such as serine (SER), cysteine (CYS), and methionine (MET), may also have slightly larger couplings involving the oxygen or sulfur atom in the side-chain. The on- site energy differences are approximately 1.0 eV for MET and CYS involving the sulfur atom, while the SER involving the oxygen atom has an on-site energy difference as large as 2.0-3.0 eV. The couplings for ILE/ADE are relatively large for the LUMO orbitals. However, their on-site energy difference is as large as 4.0 eV. Similar findings are observed with TB parameters calculated at the B3LYP level (Figure S2). Averaged over all amino acids, the nucleobases have the largest charge transfer integrals for THY (0.026 eV) and ADE (0.023 eV), followed by GUA (0.018 eV) and CYT (0.021 eV). The same trend is observed for the LUMO orbitals, where the largest charge transfer integrals have a larger value for ADE (0.054 eV) and THY (0.051 eV), and a smaller value for CYT (0.050 eV) and GUA (0.033 eV).

**Figure 4.**
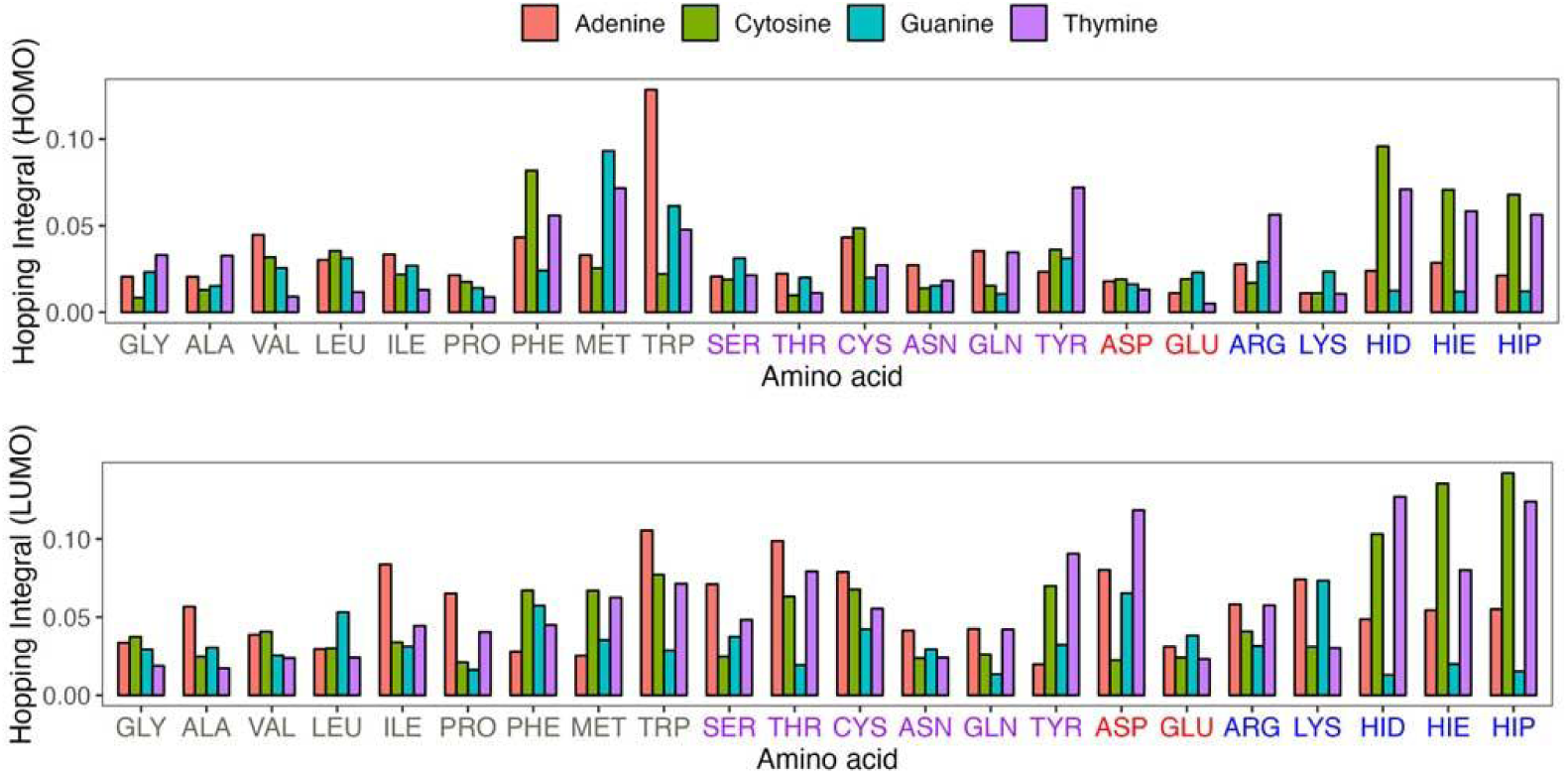
Comparative analysis of the average hopping integrals for HOMO and LUMO across AB conformers. The absolute values are used. Histidine is represented in its three protonation states: HID, HIE, and HIP. The x-axis label uses color coding to differentiate amino acids based on their chemical properties, including hydrophobicity, and polarity, acidity, basicity,

Charge transfer couplings are reported to exhibit high sensitivity to the structural orientation of molecular fragments. Figure S3 shows several AB structure contacts, where each cluster in the same AB pairs has significantly different distributions. The population of charge transfer couplings are “encoded” in various model of geometric contacts, i.e. the π-π interactions, C-H-π interactions, the hydrogen bonds or van der Waals contacts. The orientation of aromatic molecules can either enhance or diminish charge transfer couplings. The chemical diversity and specificity of various AB conformers can exhibit subtle differences in molecular structure or electronic properties, even within seemingly homogeneous groups. Note that the charge transfer couplings are not symmetric due to the inhomogeneity of DNA-protein structures, and the distribution of one type of amino acid in the frame of another reference nucleobase residue type is distinct.

As the possible structural changes will influence the electrical properties of a DNA protein complexes, the reasonable description of transfer couplings beyond the empirical formulas is very necessary. Figure 5 shows the predictive performance of the RF model for the TB parameters of arbitrary conformers, in correlating TB parameters library. The intermolecular coordinate system uses the distance (r), planar angles (θ, φ), and dihedral angles (ψ), providing a detailed set of molecular descriptors that encapsulate the spatial orientation of the molecules. The Coulomb matrix leverages atomic numbers (Z) and interatomic distances (R). This approach highlights how electronic properties are influenced by atomic identities and their spatial relationships. It emphasizes the importance of both atomic composition and geometric arrangements in determining the electronic characteristics of molecules. These descriptors are essential to machine learning models for predicting molecular properties. The correlation between actual and predicted on-site energy is very robust, with the line of best fit closely aligning with the ideal. The internal coordinates can only be successful in predicting the on-site energy, and often difficult to predict the charge transfer couplings. This suggests that the internal molecular geometries are also very important.

**Figure 5.**
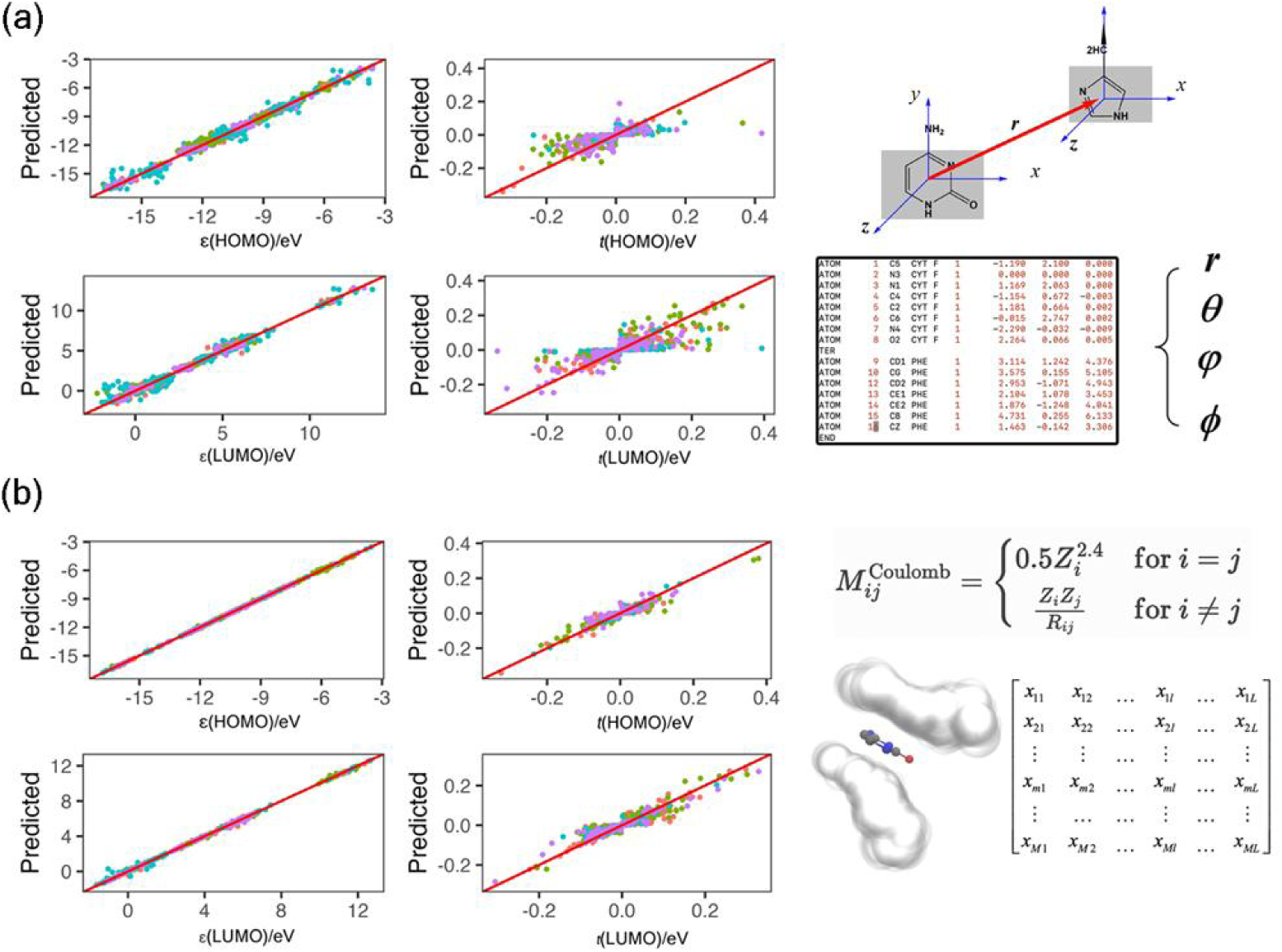
The predictive performance of the machine learning algorithm is evaluated based on two types of molecular descriptors: (a) intermolecular coordinates and (b) hydrogen-depleted Coulomb matrices. The color-coded data points represent different nucleobases

We trained the model using the 8:2 training/test ratio. Then, one could achieve a unification of accuracy and efficiency to construct TB Hamiltonian for realistic DNA, protein or DNA-protein complexes. To facilitate the use of experimental DNA and protein structures, we also compare the molecular descriptors with and without hydrogen atoms, and the results are shown in Table S1. The possibility of prediction errors in certain scenarios could lead to outliers, we have established criteria for identifying similarity between descriptors. These criteria include an average distance of less than 0.1 Å between two descriptors treated as vectors, and an angle of less than 30 degrees between multidimensional vectors exceeding three dimensions. This involves ensuring that the average distance between any two descriptors, viewed as vectors, is less than 0.1 Å, and the angle between any vectors is less than 30 degrees.

Before examining realistic systems, we first conducted an evaluation of the performance of our TB parameters. Figure S4 compares the HOMO/LUMO gap for randomly generated one thousand of dimer and trimer conformers involving nucleobases or amino acids. The results indicate that our prediction algorithm achieves deviations of 0.1∼0.2 eV, which is quite successful for such simplified TB model. The randomly generated dimers and trimers for AA configurations were derived from existing PDB databases, BB structures were partly derived from PDB and partly generated by x3DNA, while mixed AB structures were mainly derived from dimer and trimer structures at transcription factor binding sites. The insights gained from these benchmarks can be used to optimize computational strategies for modeling biological systems. In addition, the HOMO/LUMO gaps for nucleobases typically reflect their electronic properties and can vary depending on the computational method used for calculations.(147–153) Because the calculated HOMO/LUMO gap at HF level is very large (9∼10 eV) than experimental values, while B3LYP provide reasonable results (4.0∼5.0 eV). The TB parameters derived from B3LYP calculations would be used for realistic DNA-protein complexes in the following discussions.

The applicability of the BioTinter-1m model was evaluated by studying transcription factors, which are key proteins in the regulation of gene expression. They modulate the activation and repression of specific genes by binding to adjacent DNA sequences. Each transcription factor recognizes and binds to a specific sequence in the DNA alphabet (A, C, G, and T) known as a consensus site. Jun protein is an AP-1 protein, that recognizes two versions of a 7-base pair response element (Figure 6b), either TRE (5΄-TGAGTCA-3΄) with PDB ID: 2H7H or meTRE (5΄- MGAGTCA-3΄) where M = 5-methylcytosine, with PDB ID: 5T01. These elements differ only at the first base pair (bp): with T:A in TRE and 5mC:G (M:G) in meTRE. c-Jun can form both homodimers and heterodimers. Epstein-Barr Virus (EBV) Zta is a key transcription factor of the viral lytic cycle that is homologous to AP-1. The EBV viral genome is unmethylated, but becomes highly methylated during the latent stage of the viral cycle.(154, 155) Figure 6a illustrates the amino acid sequences of the human Jun protein, the Epstein-Barr virus Zta protein, and a mutant variant of the Zta protein (S186A), referred to as Zta* in this study. Zta* is designed to mimic the AP-1 protein in its interaction with the TRE DNA element, with the comparison based on the crystal structure identified by PDB ID: 2C9L. Both human AP-1 and EBV Zta are bZIP family transcription factors that bind the classical TRE. They also recognize methylated cytosine residues within different sequence contexts.(156, 157) The extensive TB parameters library is large enough to represent most possible AA, BB and AB conformers found in realistic DNA and protein structures, with prediction failures under 5% across different systems. The introduction of the BioTinter-10m model, encompassing ten million conformers, is anticipated to drastically reduce prediction errors to less than 0.1%. This process utilizes both the extensive TB parameters library and a minimal set of on-the-fly *ab initio* calculations, ensuring the robustness and accuracy of our predictions.

**Figure 6:**
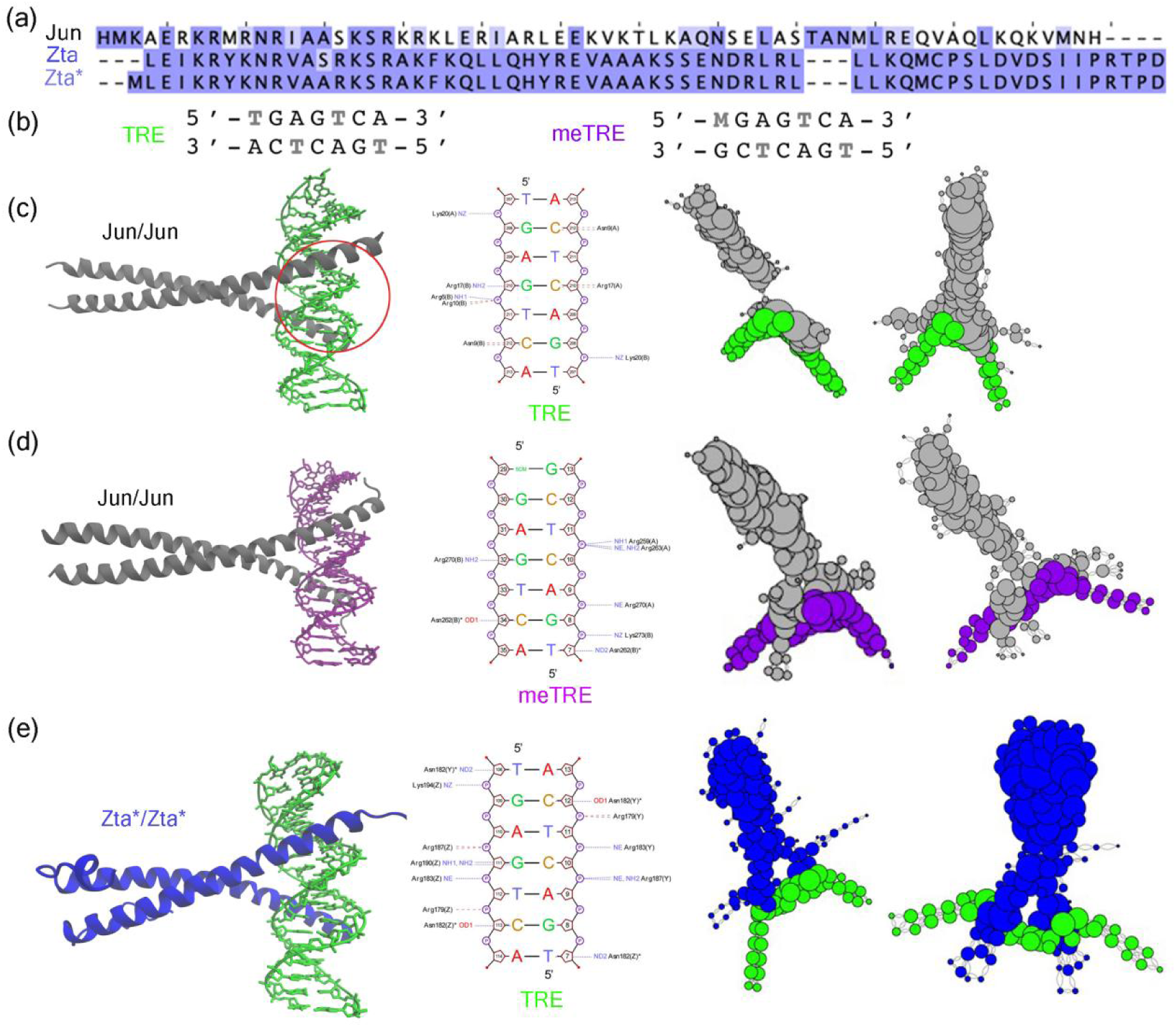
Comparative analysis of the protein-DNA interaction complexes. (a) Sequence alignment of the protein Jun, Zta and Zta* with highlighted differences. (b) Sequence alignment of the DNA elements TRE, meTRE. Visualization of (c) Jun/Jun binding to TRE, (d) Jun/Jun binding to meTRE, and (e) Zta*/Zta* binding to TRE, with the DNA-protein interface marked by a red circle, and corresponding charge transfer networks analysis for HOMO and LUMO orbitals. The size of a network node is related to its degree within the network.

Figure 6 presents a comprehensive view of the interaction between transcription factors and DNA. Each three-dimensional structure is accompanied by a schematic diagram of DNA-protein interface, highlighting the interactions between amino acids and nucleotides, and is complemented by a graphical representation of the charge transfer network. The electronic Hamiltonian of biological molecules diverges from the simple tridiagonal matrix characteristic of linear molecules due to the complex stacking arrangements of nucleobases and amino acids found in actual DNA- protein structures. In prior research, the concept of a knowledge graph was introduced as a visualization tool for TB Hamiltonian for Biomolecules. In order to construct the DNA-protein charge transfer network, each residue is represented by a vertex in the graph, and the edge represents the strength of charge transfer coupling among residues. To keep similar geometric feature as the TF molecules, we use the Kamada-Kawai layout to generate the complex network. The Kamada-Kawai algorithm is a force-directed graph layout algorithm that emphasizes the consistency between the geometric distances and graph-theoretic distances between nodes.(158) The threshold of significant charge transfer coupling is set to be 0.001 eV in this work.

Methylation can cause significant changes in DNA-protein interactions, which may result in notable alterations in gene expression patterns. Variations in nucleic acid sequences can have a significant impact on the distribution of TB Hamiltonian matrix elements at the nucleic acid- protein interface. This is demonstrated in the binding of the Jun/Jun protein to TRE and meTRE sequences, as shown in Figures 6c and 6d. Similarly, alterations in protein sequences impact both the protein termini and the nucleic acid-protein interface. This is exemplified in the interactions of Jun and the Zta* mutant protein with the TRE sequence in Figure 6c and 6e. Charge transfer networks in these DNA-protein complexes, illustrating the intricate pathways of electronic interactions within the binding interface.

After constructing the TB Hamiltonian matrix using the BioTinter-1m model for a DNA- protein complex, the direct diagonalization technique is applied to calculate various electronic structure properties. Currently, the HOMO and LUMO orbitals for each site are used as the basis functions, of course additional frontier orbitals could be easily included in our model as basis functions. As shown in Figure 7, the HOMO/LUMO gap in water solvent is larger than that in vacuum. This is quite similar as the results for model systems with DFT calculations. The frontier orbitals, especially the HOMO and LUMO orbitals are highlighted with complex network methodologies (Figure 7). The network displays the molecular orbital with larger node size for each residue that has large coefficients. The location of frontier orbitals is generally limited to a few amino acids and nucleobases. The distance between nodes is related to their sequence distance. Adjacent nodes on the network, indicate they are relatively close in secondary sequence structures.

**Figure 7.**
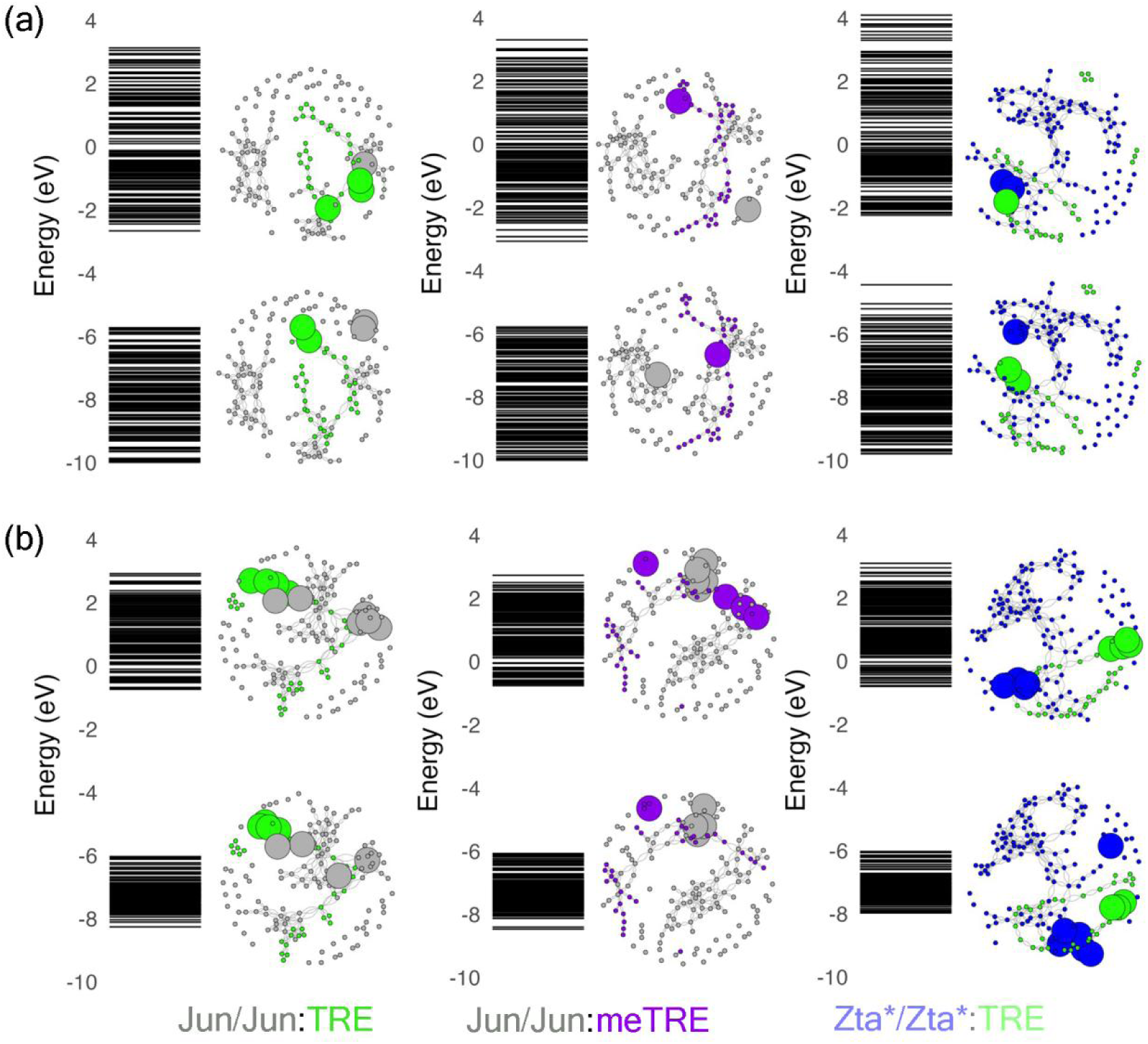
Comparative visualization of molecular orbitals across energy levels and the corresponding HOMO/LUMO distributions with complex network representation for different transcription factor-DNA complexes in (a) vacuum and (b) implicit water solvent. The coloring scheme is the same as Figure 6.

Despite its simplifications, the complex network analysis demonstrates an exceptional ability to place electronic structure variants on equal footing. The distribution of the HOMO and LUMO orbitals is generally much more dispersed in the implicit solvent model than in the vacuum model. The frontier orbitals have very distinct feature for each kind of DNA-protein complex. It is interesting to note that this structurally important residue identified as a hub is observed at the DNA-protein interface or the boundary residues of the DNA chain. In the computational model, the number of residues in the DNA chain generally does not exceed twenty residues, which may lead to boundary residues contributing to the frontier molecular orbitals. For the Jun/Jun:meTRE complex, the HOMO/LUMO orbitals are primarily distributed across amino acids and nucleobases that are relatively distant from each other. This distribution could indicate that the electronic structure of the complex facilitates charge transfer over long distances, a phenomenon that is crucial for many biological processes, such as signal transduction and energy transfer. This is consistent with the report that Methylation may cause significant changes to the photo-stability of nucleic acids, resulting in these sites becoming mutational hotspots for diseases such as skin cancer. This analysis is helpful to unravel the richness of biological electronic structure variants in realistic DNA binding protein complexes, which would evolve with fluctuating biomolecules structures.

Figure 8 presents a comparative analysis of the electronic structures of DNA-protein complexes. The analysis is presented through their density of states (DOS) under vacuum and aqueous conditions. The electronic properties are significantly influenced by solvent effects, which shift and broaden energy states around the HOMO and LUMO levels, as detailed in Figure 7. This demonstrates the role of the solvent in stabilizing electronic states. The peaks in the DOS become more pronounced and concentrated, and there are alterations in peak positions and substantial changes in peak intensities. These changes underscore the critical impact of the solvent on the electronic properties at the DNA-protein interface, where HOMO and LUMO are predominantly associated with interfacial residues. The Mulliken charges for each residue were calculated. Figure S5 displays scatter plots of the Mulliken charge populations for DNA/protein complexes in both vacuum and aqueous environments. A consistent pattern emerges across the complexes Jun/Jun:TRE, Jun/Jun:meTRE, and Zta*/Zta*:TRE, where the distribution of charges on amino acids and nucleobases appears relatively stable in water but exhibits subtle shifts in vacuum.

**Figure 8.**
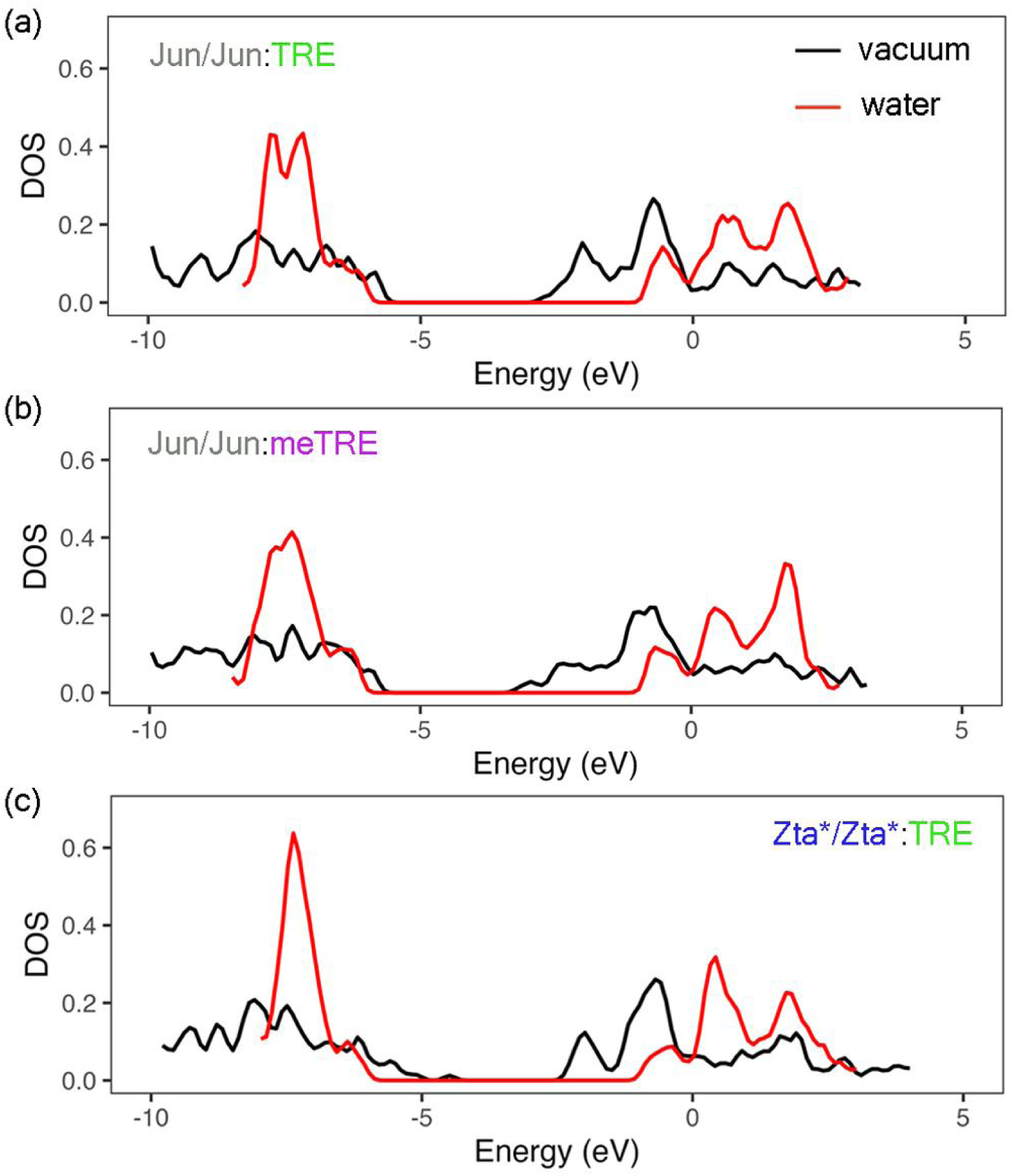
The density of states (DOS) plots for DNA-protein complexes are illustrated, contrasting calculations in vacuum (black line) with those in a water solvent environment (red line). (a) The Jun/Jun homodimer interacting with the TRE response element. (b) The Jun/Jun homodimer with the meTRE response element. (c) The Zta*/Zta* homodimer with the TRE element.

## Conclusions

Protein-DNA interactions are essential for various cellular processes such as replication, transcription, recombination, and DNA repair. Here, a library of Tight-Binding (TB) parameters has been derived for amino acids and nucleobases, containing millions of conformers. Machine learning methods were used to predict TB parameters for arbitrary fragments of amino acids and nucleobases. The electronic structure variants of the AP-1 and Epstein-Barr Virus Zta transcription factors were studied in relation to their respective transcription factor sequences and binding DNA sequences. The direct diagonalization scheme was utilized to obtain the tight- binding molecule orbitals. Our results, including DOS and frontier molecular orbitals, demonstrate significant variations in electronic structure as the protein or DNA sequence changes. This work presents a cost-effective computational tool for analyzing the electronic structure of DNA-protein structures. These insights contribute to exemplify the complex interdependence of structure, sequence, and electronic properties in the regulation of gene expression.

## Acknowledge

This work is supported by Jinzhou Medical University and Shandong University. L. Du acknowledges the support of the National Natural Science Foundation of China (No. 21503249). The authors also acknowledge the prior contributions of Prof. Chengbu Liu’s group members to the nucleic acid model systems.

## Supporting information

The details on the predictive performance of the machine learning algorithm, the TB parameters library at B3LYP level, the benchmark results, and the Mulliken charge distribution for DNA- protein complexes are given in the supporting information.

## Reference

1. Cozzolino, F., Iacobucci, I., Monaco, V. and Monti, M. (2021) Protein–DNA/RNA Interactions: An Overview of Investigation Methods in the -Omics Era. J. Proteome Res., 20, 3018–3030.

2. Luscombe, N. M. and Thornton, J. M. (2002) Protein–DNA Interactions: Amino Acid Conservation and the Effects of Mutations on Binding Specificity. J. Mol. Biol., 320, 991–1009.

3. Luscombe, N. M., Austin, S. E., Berman, H. M. and Thornton, J. M. (2000) An overview of the structures of protein-DNA complexes. Genome Biol., 1, 1–37.

4. Siggers, T. and Gordân, R. (2014) Protein–DNA binding: complexities and multi- protein codes. Nucleic Acids Res., 42, 2099–2111.

5. Grimley, E., Liao, C., Ranghini, E. J., Nikolovska-Coleska, Z. and Dressler, G. R. (2017) Inhibition of Pax2 Transcription Activation with a Small Molecule that Targets the DNA Binding Domain. ACS Chem. Biol., 12, 724–734.

6. Mitchell, P. J. and Tjian, R. (1989) Transcriptional Regulation in Mammalian Cells by Sequence-Specific DNA Binding Proteins. Science, 245, 371–378.

7. Nikolov, D. B. and Burley, S. K. (1997) RNA polymerase II transcription initiation: A structural view. Proc. Natl. Acad. Sci., 94, 15–22.

8. Lee, T. I. and Young, R. A. (2000) TRANSCRIPTION OF EUKARYOTIC PROTEIN-CODING GENES. Annu. Rev. Genet., 34, 77–137.

9. Vinson, C., Myakishev, M., Acharya, A., Mir, A. A., Moll, J. R. and Bonovich, M. (2002) Classification of Human B-ZIP Proteins Based on Dimerization Properties. Mol. Cell. Biol., 22, 6321–6335.

10. Babu, M. M., Luscombe, N. M., Aravind, L., Gerstein, M. and Teichmann, S. A. (2004) Structure and evolution of transcriptional regulatory networks. Curr. Opin. Struct. Biol., 14, 283–291.

11. Lin, M. and Guo, J. (2019) New insights into protein–DNA binding specificity from hydrogen bond based comparative study. Nucleic Acids Res., 47, 11103–11113.

12. Olins, A. L. and Olins, D. E. (1974) Spheroid Chromatin Units (ν Bodies). Science, 183, 330–332.

13. Kornberg, R. D. and Thomas, J. O. (1974) Chromatin Structure: Oligomers of the Histones. Science, 184, 865–868.

14. Kornberg, R. D. (1974) Chromatin Structure: A Repeating Unit of Histones and DNA. Science, 184, 868–871.

15. Fyodorov, D. V., Zhou, B. -R., Skoultchi, A. I. and Bai, Y. (2018) Emerging roles of linker histones in regulating chromatin structure and function. Nat. Rev. Mol. Cell Biol., 19, 192–206.

16. Luger, K., Rechsteiner, T. J., Flaus, A. J., Waye, M. M. Y. and Richmond, T. J. (1997) Characterization of nucleosome core particles containing histone proteins made in bacteria1. J. Mol. Biol., 272, 301–311.

17. Schlissel, G. and Rine, J. (2019) The nucleosome core particle remembers its position through DNA replication and RNA transcription. Proc. Natl. Acad. Sci., 116, 20605–20611.

18. Sato, S., Takizawa, Y., Hoshikawa, F., Dacher, M., Tanaka, H., Tachiwana, H., Kujirai, T., Iikura, Y., Ho, C. -H., Adachi, N., et al. (2021) Cryo-EM structure of the nucleosome core particle containing Giardia lamblia histones. Nucleic Acids Res., 49, 8934–8946.

19. Latchman, D. S. (1997) Transcription factors: An overview. Int. J. Biochem. Cell Biol., 29, 1305–1312.

20. Ptashne, M. and Gann, A. (1997) Transcriptional activation by recruitment. Nature, 386, 569–577.

21. Karin, M. (1990) Too many transcription factors: positive and negative interactions. New Biol., 2, 126–131.

22. Nakabeppu, Y., Ryder, K. and Nathans, D. (1988) DNA binding activities of three murine Jun proteins: Stimulation by Fos. Cell, 55, 907–915.

23. Rauscher, F. J., Voulalas, P. J., Franza, B. R. and Curran, T. (1988) Fos and Jun bind cooperatively to the AP-1 site: reconstitution in vitro. Genes Dev., 2, 1687–1699.

24. Kataoka, K., Noda, M. and Nishizawa, M. (1994) Maf nuclear oncoprotein recognizes sequences related to an AP-1 site and forms heterodimers with both Fos and Jun. Mol. Cell. Biol., 14, 700–712.

25. Hess, J., Angel, P. and Schorpp-Kistner, M. (2004) AP-1 subunits: quarrel and harmony among siblings. J. Cell Sci., 117, 5965–5973.

26. Hai, T. and Curran, T. (1991) Cross-family dimerization of transcription factors Fos/Jun and ATF/CREB alters DNA binding specificity. Proc. Natl. Acad. Sci., 88, 3720–3724.

27. Karin, M., Liu, Z. and Zandi, E. (1997) AP-1 function and regulation. Curr. Opin. Cell Biol., 9, 240–246.

28. Jakoby, M., Weisshaar, B., Dröge-Laser, W., Vicente-Carbajosa, J., Tiedemann, J., Kroj, T. and Parcy, F. (2002) bZIP transcription factors in Arabidopsis. Trends Plant Sci., 7, 106–111.

29. Eferl, R. and Wagner, E. F. (2003) AP-1: a double-edged sword in tumorigenesis. Nat. Rev. Cancer, 3, 859–868.

30. Liao, Y., Zou, H. -F., Wang, H. -W., Zhang, W. -K., Ma, B., Zhang, J. -S. and Chen, S. -Y. (2008) Soybean GmMYB76, GmMYB92, and GmMYB177 genes confer stress tolerance in transgenic Arabidopsis plants. *Cell Res.*, **18**, 1047–1060.

31. Uluçkan, Ö., Guinea-Viniegra, J., Jimenez, M. and Wagner, E. F. (2015) Signalling in inflammatory skin disease by AP-1 (Fos/Jun). Clin. Exp. Rheumatol.

32. Papoudou-Bai, A., Hatzimichael, E., Barbouti, A. and Kanavaros, P. (2017) Expression patterns of the activator protein-1 (AP-1) family members in lymphoid neoplasms. Clin. Exp. Med., 17, 291–304.

33. Martínez-Zamudio, R. I., Roux, P. -F., de Freitas, J. A. N. L. F., Robinson, L., Doré, G., Sun, B., Belenki, D., Milanovic, M., Herbig, U., Schmitt, C. A., et al. (2020) AP-1 imprints a reversible transcriptional programme of senescent cells. Nat. Cell Biol., 22, 842–855.

34. Wu, Z., Nicoll, M. and Ingham, R. J. (2021) AP-1 family transcription factors: a diverse family of proteins that regulate varied cellular activities in classical hodgkin lymphoma and ALK+ ALCL. Exp. Hematol. Oncol., 10, 4.

35. Schreiber, M., Kolbus, A., Piu, F., Szabowski, A., Möhle-Steinlein, U., Tian, J., Karin, M., Angel, P. and Wagner, E. F. (1999) Control of cell cycle progression by c- Jun is p53 dependent. Genes Dev., 13, 607–619.

36. Shaulian, E. (2010) AP-1 — The Jun proteins: Oncogenes or tumor suppressors in disguise? Cell. Signal., 22, 894–899.

37. Sengupta, S., Ghufran, S. M., Khan, A., Biswas, S. and Roychoudhury, S. (2022) Transition of amyloid/mutant p53 from tumor suppressor to an oncogene and therapeutic approaches to ameliorate metastasis and cancer stemness. Cancer Cell Int., 22, 416.

38. Bahar, M. E., Kim, H. J. and Kim, D. R. (2023) Targeting the RAS/RAF/MAPK pathway for cancer therapy: from mechanism to clinical studies. Signal Transduct. Target. Ther., 8, 455.

39. Mondol, T., Batabyal, S. and Pal, S. K. (2012) Ultrafast electron transfer in the recognition of different DNA sequences by a DNA-binding protein with different dynamical conformations. J. Biomol. Struct. Dyn., 30, 362–370.

40. Batabyal, S., Choudhury, S., Sao, D., Mondol, T. and Pal, S. K. (2014) Dynamical perspective of protein-DNA interaction. Biomol. Concepts, 5, 21–43.

41. Choudhury, S., Naiya, G., Singh, P., Lemmens, P., Roy, S. and Pal, S. K. (2016) Modulation of Ultrafast Conformational Dynamics in Allosteric Interaction of Gal Repressor Protein with Different Operator DNA Sequences. ChemBioChem, 17, 605– 613.

42. Choudhury, S., Naiya, G., Singh, P., Lemmens, P., Roy, S. and Pal, S. K. (2016) Inside Cover: Modulation of Ultrafast Conformational Dynamics in Allosteric Interaction of Gal Repressor Protein with Different Operator DNA Sequences (ChemBioChem 7/2016). ChemBioChem, 17, 524–524.

43. Choudhury, S., Ghosh, B., Singh, P., Ghosh, R., Roy, S. and Pal, S. K. (2016) Ultrafast differential flexibility of Cro-protein binding domains of two operator DNAs with different sequences. Phys. Chem. Chem. Phys., 18, 17983–17990.

44. Cellini, A., Shankar, M. K., Nimmrich, A., Hunt, L. A., Monrroy, L., Mutisya, J., Furrer, A., Beale, E. V., Carrillo, M., Malla, T. N., et al. (2024) Directed ultrafast conformational changes accompany electron transfer in a photolyase as resolved by serial crystallography. Nat. Chem., 10. 1038/s41557-023-01413-9.

45. Hall, D. B., Holmlin, R. E. and Barton, J. K. (1996) Oxidative DNA damage through long-range electron transfer. Nature, 382, 731–735.

46. Carell, T., Burgdorf, L. T., Kundu, L. M. and Cichon, M. (2001) The mechanism of action of DNA photolyases. Curr. Opin. Chem. Biol., 5, 491–498.

47. Rogozin, I. B. and Pavlov, Y. I. (2003) Theoretical analysis of mutation hotspots and their DNA sequence context specificity. Mutat. Res. Mutat. Res., 544, 65–85.

48. Delaney, S. and Barton, J. K. (2003) Long-Range DNA Charge Transport. J. Org. Chem., 68, 6475–6483.

49. Tashiro, R., Wang, A. H. -J. and Sugiyama, H. (2006) Photoreactivation of DNA by an Archaeal nucleoprotein Sso7d. Proc. Natl. Acad. Sci., 103, 16655–16659.

50. Hatcher, E., Balaeff, A., Keinan, S., Venkatramani, R. and Beratan, D. N. (2008) PNA versus DNA: Effects of Structural Fluctuations on Electronic Structure and Hole-Transport Mechanisms. J. Am. Chem. Soc., 130, 11752–11761.

51. Boal, A. K., Genereux, J. C., Sontz, P. A., Gralnick, J. A., Newman, D. K. and Barton, J. K. (2009) Redox signaling between DNA repair proteins for efficient lesion detection. Proc. Natl. Acad. Sci., 106, 15237–15242.

52. Morinaga, H., Takenaka, T., Hashiya, F., Kizaki, S., Hashiya, K., Bando, T. and Sugiyama, H. (2013) Sequence-specific electron injection into DNA from an intermolecular electron donor. Nucleic Acids Res., 41, 4724–4728.

53. Beratan, D. N. (2019) Why Are DNA and Protein Electron Transfer So Different? Annu. Rev. Phys. Chem., 70, 71–97.

54. Hashiya, F., Ito, S. and Sugiyama, H. (2019) Electron injection from mitochondrial transcription factor A to DNA associated with thymine dimer photo repair. Bioorg. Med. Chem., 27, 278–284.

55. Ladik, J., Bende, A. and Bogár, F. (2010) Charge transfer between DNA and proteins in the nucleosomes. Theor. Chem. Acc., 125, 185–191.

56. Sontz, P. A., Mui, T. P., Fuss, J. O., Tainer, J. A. and Barton, J. K. (2012) DNA charge transport as a first step in coordinating the detection of lesions by repair proteins. Proc. Natl. Acad. Sci., 109, 1856–1861.

57. Grodick, M. A., Muren, N. B. and Barton, J. K. (2015) DNA Charge Transport within the Cell. Biochemistry, 54, 962–973.

58. Arnold, A. R., Grodick, M. A. and Barton, J. K. (2016) DNA Charge Transport: from Chemical Principles to the Cell. Cell Chem. Biol., 23, 183–197.

59. Fuss, J. O., Tsai, C. -L., Ishida, J. P. and Tainer, J. A. (2015) Emerging critical roles of Fe–S clusters in DNA replication and repair. Biochim. Biophys. Acta BBA - Mol. Cell Res., 1853, 1253–1271.

60. O’Brien, E., Holt, M. E., Thompson, M. K., Salay, L. E., Ehlinger, A. C., Chazin, W. J. and Barton, J. K. (2017) The [4Fe4S] cluster of human DNA primase functions as a redox switch using DNA charge transport. Science, 355, eaag1789.

61. Tse, E. C. M., Zwang, T. J. and Barton, J. K. (2017) The Oxidation State of [4Fe4S] Clusters Modulates the DNA-Binding Affinity of DNA Repair Proteins. J. Am. Chem. Soc., 139, 12784–12792.

62. Syed, A. and Tainer, J. A. (2019) Charge Transport Communication through DNA by Protein Fe–S Clusters: How Far Is Not Too Far? ACS Cent. Sci., 5, 7–9.

63. Derr, J. B., Tamayo, J., Clark, J. A., Morales, M., Mayther, M. F., Espinoza, E. M., Rybicka-Jasińska, K. and Vullev, V. I. (2020) Multifaceted aspects of charge transfer. Phys. Chem. Chem. Phys., 22, 21583–21629.

64. Fox, S. J., Dziedzic, J., Fox, T., Tautermann, C. S. and Skylaris, C. -K. (2014) Density functional theory calculations on entire proteins for free energies of binding: Application to a model polar binding site. Proteins Struct. Funct. Bioinforma., 82, 3335–3346.

65. He, X., Zhu, T., Wang, X., Liu, J. and Zhang, J. Z. H. (2014) Fragment Quantum Mechanical Calculation of Proteins and Its Applications. Acc. Chem. Res., 47, 2748– 2757.

66. Koch, T., Shim, I., Lindow, M., Ørum, H. and Bohr, H. G. (2014) Quantum Mechanical Studies of DNA and LNA. Nucleic Acid Ther., 24, 139–148.

67. Deng, A., Li, H., Bo, M., Huang, Z., Li, L., Yao, C. and Li, F. (2020) Understanding atomic bonding and electronic distributions of a DNA molecule using DFT calculation and BOLS-BC model. Biochem. Biophys. Rep., 24, 100804.

68. Gundelach, L., Fox, T., S. Tautermann, C. and Skylaris, C. -K. (2021) Protein–ligand free energies of binding from full-protein DFT calculations: convergence and choice of exchange–correlation functional. Phys. Chem. Chem. Phys., 23, 9381–9393.

69. Slater, J. C. and Koster, G. F. (1954) Simplified LCAO Method for the Periodic Potential Problem. Phys. Rev., 94, 1498–1524.

70. Goringe, C. M., Bowler, D. R. and Hernández, E. (1997) Tight-binding modelling of materials. Rep. Prog. Phys., 60, 1447.

71. Conwell, E. M. and Rakhmanova, S. V. (2000) Polarons in DNA. Proc. Natl. Acad. Sci., 97, 4556–4560.

72. Grimme, S., Bannwarth, C. and Shushkov, P. (2017) A Robust and Accurate Tight- Binding Quantum Chemical Method for Structures, Vibrational Frequencies, and Noncovalent Interactions of Large Molecular Systems Parametrized for All spd-Block Elements (Z = 1–86). *J. Chem. Theory Comput.*, **13**, 1989–2009.

73. Spiegelman, F., Tarrat, N., Cuny, J., Dontot, L., Posenitskiy, E., Martí, C., Simon, A. and Rapacioli, M. (2020) Density-functional tight-binding: basic concepts and applications to molecules and clusters. Adv. Phys. X, 5, 1710252.

74. Vishal **dal, Janik, M. J. and Milner, S. T. (2024) Tight-binding model describes frontier orbitals of non-fullerene acceptors. Mol. Syst. Des. Eng., 9, 382–398.

75. Koslowski, T. (1999) Localized and extended electronic eigenstates in proteins: A tight-binding approach. J. Chem. Phys., 110, 12233–12239.

76. Song, J., Zhang, D. C., Liu, D. S., Mei, L. M. and Xie, S. J. (2005) Density of states of DNA molecules with varied itinerant electrons. Synth. Met., 155, 607–610.

77. Triberis, G. P. and Dimakogianni, M. (2008) Correlated small polaron hopping transport in 1D disordered systems at high temperatures: a possible charge transport mechanism in DNA. J. Phys. Condens. Matter, 21, 035114.

78. Yamada, H. and Iguchi, K. (2010) Some Effective Tight-Binding Models for Electrons in DNA Conduction: A Review. *Adv*. Condens. Matter Phys., 2010, 380710.

79. Zilly, M., Ujsághy, O. and Wolf, D. E. (2010) Conductance of DNA molecules: Effects of decoherence and bonding. Phys. Rev. B, 82, 125125.

80. Malakooti, S., Hedin, E. and Joe, Y. (2013) Tight-binding approach to strain- dependent DNA electronics. J. Appl. Phys., 114, 014701.

81. Hu, F., He, F. and Yaron, D. J. (2023) Treating Semiempirical Hamiltonians as Flexible Machine Learning Models Yields Accurate and Interpretable Results. J. Chem. Theory Comput., 19, 6185–6196.

82. Gillet, N., Berstis, L., Wu, X., Gajdos, F., Heck, A., de la Lande, A., Blumberger, J. and Elstner, M. (2016) Electronic Coupling Calculations for Bridge-Mediated Charge Transfer Using Constrained Density Functional Theory (CDFT) and Effective Hamiltonian Approaches at the Density Functional Theory (DFT) and Fragment- Orbital Density Functional Tight Binding (FODFTB) Level. J. Chem. Theory Comput., 12, 4793–4805.

83. Beratan, D. N., Onuchic, J. N. and Hopfield, J. J. (1987) Electron tunneling through covalent and noncovalent pathways in proteins. J. Chem. Phys., 86, 4488–4498.

84. Grozema, F. C., Berlin, Y. A. and Siebbeles, L. D. A. (2000) Mechanism of Charge Migration through DNA: Molecular Wire Behavior, Single-Step Tunneling or Hopping? J. Am. Chem. Soc., 122, 10903–10909.

85. Balabin, I. A. and Onuchic, J. N. (2000) Dynamically Controlled Protein Tunneling Paths in Photosynthetic Reaction Centers. Science, 290, 114–117.

86. de la Lande, A. and Salahub, D. R. (2010) Derivation of interpretative models for long range electron transfer from constrained density functional theory. J. Mol. Struct. THEOCHEM, 943, 115–120.

87. de la Lande, A., Babcock, N. S., Řezáč, J., Sanders, B. C. and Salahub, D. R. (2010) Surface residues dynamically organize water bridges to enhance electron transfer between proteins. Proc. Natl. Acad. Sci., 107, 11799–11804.

88. Balabin, I. A., Hu, X. and Beratan, D. N. (2012) Exploring biological electron transfer pathway dynamics with the Pathways Plugin for VMD. J. Comput. Chem., 33, 906–910.

89. Hammi, E. E., Houée-Lévin, C., Řezáč, J., Lévy, B., Demachy, I., Baciou, L. and Lande, A. de la (2012) New insights into the mechanism of electron transfer within flavohemoglobins: tunnelling pathways, packing density, thermodynamic and kinetic analyses. Phys. Chem. Chem. Phys., 14, 13872–13880.

90. Mcmahan, A. K. and Klepeis, J. E. (1997) Ab Initio Calculation of Tight-Binding Parameters. MRS Proc., 491, 199.

91. Agapito, L. A., Ismail-Beigi, S., Curtarolo, S., Fornari, M. and Nardelli, M. B. (2016) Accurate tight-binding Hamiltonian matrices from ab initio calculations: Minimal basis sets. Phys. Rev. B, 93, 035104.

92. Wang, H., Liu, F., Dong, T., Du, L., Zhang, D. and Gao, J. (2018) Charge-Transfer Knowledge Graph among Amino Acids Derived from High-Throughput Electronic Structure Calculations for Protein Database. ACS Omega, 3, 4094–4104.

93. Berman, H. M., Westbrook, J., Feng, Z., Gilliland, G., Bhat, T. N., Weissig, H., Shindyalov, I. N. and Bourne, P. E. (2000) The Protein Data Bank. Nucleic Acids Res., 28, 235–242.

94. Lu, X. and Olson, W. K. (2003) 3DNA: a software package for the analysis, rebuilding and visualization of three-dimensional nucleic acid structures. Nucleic Acids Res., 31, 5108–5121.

95. Norambuena, T. and Melo, F. (2010) The Protein-DNA Interface database. BMC Bioinformatics, 11, 1–12.

96. Baek, M., McHugh, R., Anishchenko, I., Jiang, H., Baker, D. and DiMaio, F. (2024) Accurate prediction of protein–nucleic acid complexes using RoseTTAFoldNA. Nat. Methods, 21, 117–121.

97. Liu, F. and Du, L. (2023) The Charge Transfer Network Model for Arbitrary Proteins Complexes. In Wen, S., Yang, C. (eds), Biomedical and Computational Biology. Springer International Publishing, Cham, pp. 1–12.

98. Behler, J. and Parrinello, M. (2007) Generalized Neural-Network Representation of High-Dimensional Potential-Energy Surfaces. Phys. Rev. Lett., 98, 146401.

99. Braams, B. J. and Bowman, J. M. (2009) Permutationally invariant potential energy surfaces in high dimensionality. Int. Rev. Phys. Chem., 28, 577–606.

100. Bartók, A. P., Payne, M. C., Kondor, R. and Csányi, G. (2010) Gaussian Approximation Potentials: The Accuracy of Quantum Mechanics, without the Electrons. Phys. Rev. Lett., 104, 136403.

101. Smith, J. S., Isayev, O. and Roitberg, A. E. (2017) ANI-1: an extensible neural network potential with DFT accuracy at force field computational cost. Chem. Sci., 8, 3192–3203.

102. Podryabinkin, E. V. and Shapeev, A. V. (2017) Active learning of linearly parametrized interatomic potentials. Comput. Mater. Sci., 140, 171–180.

103. Keith, J. A., Vassilev-Galindo, V., Cheng, B., Chmiela, S., Gastegger, M., Müller, K. -R. and Tkatchenko, A. (2021) Combining Machine Learning and Computational Chemistry for Predictive Insights Into Chemical Systems. Chem. Rev., 121, 9816–9872.

104. Hagg, A. and Kirschner, K. N. (2023) Open-Source Machine Learning in Computational Chemistry. J. Chem. Inf. Model., 63, 4505–4532.

105. Dral, P. O. (2024) AI in computational chemistry through the lens of a decade-long journey. Chem. Commun., 60, 3240–3258.

106. Back, S., Aspuru-Guzik, A., Ceriotti, M., Gryn’ova, G., Grzybowski, B., Gu, G. H., Hein, J., Hippalgaonkar, K., Hormázabal, R., Jung, Y., et al. (2024) Accelerated chemical science with AI. Digit. Discov., 3, 23–33.

107. Chmiela, S., Sauceda, H. E., Müller, K. -R. and Tkatchenko, A. (2018) Towards exact molecular dynamics simulations with machine-learned force fields. Nat. Commun., 9, 3887.

108. Chmiela, S., Tkatchenko, A., Sauceda, H. E., Poltavsky, I., Schütt, K. T. and Müller, K. -R. (2017) Machine learning of accurate energy-conserving molecular force fields. Sci. Adv., 3, e1603015.

109. Ryczko, K., Strubbe, D. A. and Tamblyn, I. (2019) Deep learning and density-functional theory. Phys. Rev. A, 100, 022512.

110. Brockherde, F., Vogt, L., Li, L., Tuckerman, M. E., Burke, K. and Müller, K. -R. (2017) Bypassing the Kohn-Sham equations with machine learning. Nat. Commun., 8, 872.

111. Guzman-Pando, A., Ramirez-Alonso, G., Arzate-Quintana, C. and Camarillo- Cisneros, J. (2023) Deep learning algorithms applied to computational chemistry. Mol. Divers., 10. 1007/s11030-023-10771-y.

112. Wilkins, D. M., Grisafi, A., Yang, Y., Lao, K. U., DiStasio, R. A. and Ceriotti, M. (2019) Accurate molecular polarizabilities with coupled cluster theory and machine learning. Proc. Natl. Acad. Sci., 116, 3401–3406.

113. Gastegger, M., Behler, J. and Marquetand, P. (2017) Machine learning molecular dynamics for the simulation of infrared spectra. Chem. Sci., 8, 6924–6935.

114. Schütt, K. T., Gastegger, M., Tkatchenko, A., Müller, K. -R. and Maurer, R. J. (2019) Unifying machine learning and quantum chemistry with a deep neural network for molecular wavefunctions. Nat. Commun., 10, 5024.

115. Sugden, B. (2014) Epstein-Barr Virus: The Path from Association to Causality for a Ubiquitous Human Pathogen. PLOS Biol., 12, e1001939.

116. Chiu, Y. -F. and Sugden, B. (2016) Epstein-Barr Virus: The Path from Latent to Productive Infection. Annu. Rev. Virol., 3, 359–372.

117. Cui, P., Wu, J., Zhang, G. and Liu, C. (2008) Hole polarons in poly(G)-poly(C) and poly(A)-poly(T) DNA molecules. Sci. China Ser. B Chem., 51, 1182–1186.

118. Luscombe, N. M., Laskowski, R. A. and Thornton, J. M. (2001) Amino acid–base interactions: a three-dimensional analysis of protein–DNA interactions at an atomic level. Nucleic Acids Res., 29, 2860–2874.

119. Case, D. A., Aktulga, H. M., Belfon, K., Cerutti, D. S., Cisneros, G. A., Cruzeiro, V. W. D., Forouzesh, N., Giese, T. J., Götz, A. W., Gohlke, H., et al. (2023) AmberTools. J. Chem. Inf. Model., 63, 6183–6191.

120. Zheng, B., Wu, J., Sun, W. and Liu, C. (2006) Trapping and hopping of polaron in DNA periodic stack. Chem. Phys. Lett., 425, 123–127.

121. Canola, S., Pecoraro, C. and Negri, F. (2016) Dimer and cluster approach for the evaluation of electronic couplings governing charge transport: Application to two pentacene polymorphs. Chem. Phys., 478, 130–138.

122. Valeev, E. F., Coropceanu, V., da Silva Filho, D. A., Salman, S. and Brédas, J. -L. (2006) Effect of Electronic Polarization on Charge-Transport Parameters in Molecular Organic Semiconductors. J. Am. Chem. Soc., 128, 9882–9886.

123. Cui, P., Zhang, D., Liu, Y., Yuan, S., Li, B., Gao, J. and Liu, C. (2011) Tight-binding model method and its applications in DNA molecules. Sci. Sin. Chim., 41, 748–755.

124. Kensert, A., Alvarsson, J., Norinder, U. and Spjuth, O. (2018) Evaluating parameters for ligand-based modeling with random forest on sparse data sets. J. Cheminformatics, 10, 49.

125. Breskvar, M., Kocev, D. and Džeroski, S. (2018) Ensembles for multi-target regression with random output selections. Mach. Learn., 107, 1673–1709.

126. Haghighatlari, M., Li, J., Heidar-Zadeh, F., Liu, Y., Guan, X. and Head-Gordon, T. (2020) Learning to Make Chemical Predictions: The Interplay of Feature Representation, Data, and Machine Learning Methods. Chem, 6, 1527–1542.

127. Sun, L., Ji, Y., Zhu, X. and Peng, T. (2022) Process knowledge-based random forest regression for model predictive control on a nonlinear production process with multiple working conditions. Adv. Eng. Inform., 52, 101561.

128. Schmid, L., Gerharz, A., Groll, A. and Pauly, M. (2023) Tree-based ensembles for multi-output regression: Comparing multivariate approaches with separate univariate ones. Comput. Stat. Data Anal., 179, 107628.

129. Mahesh R N, U. and Nelleri, A. (2023) Multi-Class Classification and Multi- Output Regression of Three-Dimensional Objects Using Artificial Intelligence Applied to Digital Holographic Information. Sensors, 23, 1095.

130. Pedregosa, F., Varoquaux, G., Gramfort, A., Michel, V., Thirion, B., Grisel, O., Blondel, M., Prettenhofer, P., Weiss, R., Dubourg, V., et al. (2011) Scikit-learn: Machine Learning in Python. J. Mach. Learn. Res., 12, 2825–2830.

131. Mitchell, J. B. O. (2014) Machine learning methods in chemoinformatics. WIREs Comput. Mol. Sci., 4, 468–481.

132. Goh, G. B., Hodas, N. O. and Vishnu, A. (2017) Deep learning for computational chemistry. J. Comput. Chem., 38, 1291–1307.

133. Korshunova, M., Ginsburg, B., Tropsha, A. and Isayev, O. (2021) OpenChem: A Deep Learning Toolkit for Computational Chemistry and Drug Design. J. Chem. Inf. Model., 61, 7–13.

134. Jeyachitra, R. K. and Manochandar, S. (2023) Machine Learning and Deep Learning. In Multimodal Biometric and Machine Learning Technologies. John Wiley & Sons, Ltd, pp. 173–225.

135. Guzman-Pando, A., Ramirez-Alonso, G., Arzate-Quintana, C. and Camarillo- Cisneros, J. (2023) Deep learning algorithms applied to computational chemistry. Mol. Divers., 10. 1007/s11030-023-10771-y.

136. James, T. and Hristozov, D. (2022) Deep Learning and Computational Chemistry. In *Artificial Intelligence in Drug Design*. Humana, New York, NY, pp. 125–151.

137. Paszke, A., Gross, S., Massa, F., Lerer, A., Bradbury, J., Chanan, G., Killeen, T., Lin, Z., Gimelshein, N., Antiga, L., et al. (2019) PyTorch: An Imperative Style, High- Performance Deep Learning Library. In Advances in Neural Information Processing Systems. Vol. 32.

138. Neese, F., Wennmohs, F., Becker, U. and Riplinger, C. (2020) The ORCA quantum chemistry program package. J. Chem. Phys., 152, 224108.

139. Frisch, M. J., Trucks, G. W., Cheeseman, J. R., Scalmani, G., Caricato, M., Hratchian, H. P., Li, X., Barone, V., Bloino, J., Zheng, G., et al. (2009) Gaussian 09.

140. Jumper, J., Evans, R., Pritzel, A., Green, T., Figurnov, M., Ronneberger, O., Tunyasuvunakool, K., Bates, R., Žídek, A., Potapenko, A., et al. (2021) Highly accurate protein structure prediction with AlphaFold. Nature, 596, 583–589.

141. Jäger, M. O. J., Morooka, E. V., Federici Canova, F., Himanen, L. and Foster, A. S. (2018) Machine learning hydrogen adsorption on nanoclusters through structural descriptors. Npj Comput. Mater., 4, 37.

142. 142. Rupp, M., Tkatchenko, A., Müller, K. -R. and von Lilienfeld, O. A. (2012) Fast and Accurate Modeling of Molecular Atomization Energies with Machine Learning. Phys. Rev. Lett., 108, 058301.

143. De, S., Bartók, A. P., Csányi, G. and Ceriotti, M. (2016) Comparing molecules and solids across structural and alchemical space. Phys. Chem. Chem. Phys., 18, 13754– 13769.

144. Willatt, M. J., Musil, F. and Ceriotti, M. (2018) Feature optimization for atomistic machine learning yields a data-driven construction of the periodic table of the elements. Phys. Chem. Chem. Phys., 20, 29661–29668.

145. Mölder, F., Jablonski, K. P., Letcher, B., Hall, M. B., Tomkins-Tinch, C. H., Sochat, V., Forster, J., Lee, S., Twardziok, S. O., Kanitz, A., et al. (2021) Sustainable data analysis with Snakemake. F1000Research, 10, 33.

146. Liu, F. and Du, L. (2024) Decoding dominant interaction patterns in halogenated dimers: A journey from halogen bonding to Van der Waals interactions. Comput. Theor. Chem., 1233, 114513.

147. Clark, L. B., Peschel, G. G. and Tinoco, I. Jr. (1965) Vapor Spectra and Heats of Vaporization of Some Purine and Pyrimidine Bases1. J. Phys. Chem., 69, 3615–3618.

148. Li, Liang. and Lubman, D. M. (1987) Ultraviolet-visible absorption spectra of biological molecules in the gas phase using pulsed laser-induced volatilization enhancement in a diode array spectrophotometer. Anal. Chem., 59, 2538–2541.

149. Ladik, J., Bende, A. and Bogár, F. (2008) The electronic structure of the four nucleotide bases in DNA, of their stacks, and of their homopolynucleotides in the absence and presence of water. J. Chem. Phys., 128, 105101.

150. Miyahara, T. and Nakatsuji, H. (2011) Absorption spectra of nucleic acid bases studied by the symmetry-adapted-cluster configuration-interaction (SAC-CI) method. Collect. Czechoslov. Chem. Commun., 76, 537–552.

151. Foster, M. E. and Wong, B. M. (2012) Nonempirically Tuned Range-Separated DFT Accurately Predicts Both Fundamental and Excitation Gaps in DNA and RNA Nucleobases. J. Chem. Theory Comput., 8, 2682–2687.

152. Leal, L. A. E. and Lopez-Acevedo, O. (2015) On the interaction between gold and silver metal atoms and DNA/RNA nucleobases – a comprehensive computational study of ground state properties. Nanotechnol. Rev., 4, 173–191.

153. Üngördü, A. and Tezer, N. (2017) The solvent (water) and metal effects on HOMO-LUMO gaps of guanine base pair: A computational study. J. Mol. Graph. Model., 74, 265–272.

154. Kenney, S. C. and Mertz, J. E. (2014) Regulation of the latent-lytic switch in Epstein–Barr virus. Semin. Cancer Biol., 26, 60–68.

155. Paulson, E. J. and Speck, S. H. (1999) Differential Methylation of Epstein-Barr Virus Latency Promoters Facilitates Viral Persistence in Healthy Seropositive Individuals. J. Virol., 73, 9959–9968.

156. Petosa, C., Morand, P., Baudin, F., Moulin, M., Artero, J. -B. and Müller, C. W. (2006) Structural Basis of Lytic Cycle Activation by the Epstein-Barr Virus ZEBRA Protein. Mol. Cell, 21, 565–572.

157. Glover, J. N. M. and Harrison, S. C. (1995) Crystal structure of the heterodimeric bZIP transcription factor c-Fos–c-Jun bound to DNA. Nature, 373, 257–261.

158. Schreiber, F., Dwyer, T., Marriott, K. and Wybrow, M. (2009) A generic algorithm for layout of biological networks. BMC Bioinformatics, 10, 1–12.

