## Supporting Information File for "Elucidating Electronic Structure Variations in Nucleic Acid-Protein Complexes Involved in Transcription Regulation Using a Tight-Binding Approach"

**Table S1**. The predictive performance of the machine learning algorithm is evaluated based various molecular descriptors. The internal coordinates (IC), Coulomb matrices (CM), and SOAP model. We also compare the molecular descriptors with and without hydrogen atoms. The correlation coefficients were summarized.

| | IC | CM | CM hydrogen-depleted | SOAP | SOAP hydrogen-depleted |
|---|---|---|---|---|---|
| HF/6-31G(d) | | | | | |
| ε(HOMO) | 0.9948 | 0.9998 | 0.9998 | 0.9999 | 0.9998 |
| ε(LUMO) | 0.9937 | 0.9997 | 0.9996 | 0.9994 | 0.9993 |
| t(HOMO) | 0.7642 | 0.9539 | 0.9579 | 0.9545 | 0.9511 |
| t(LUMO) | 0.7909 | 0.9399 | 0.9338 | 0.9408 | 0.9295 |
| B3LYP/6-31G(d) | | | | | |
| ε(HOMO) | 0.9947 | 0.9999 | 0.9999 | 0.9999 | 0.9998 |
| ε(LUMO) | 0.9949 | 0.9999 | 0.9998 | 0.9998 | 0.9997 |
| t(HOMO) | 0.8154 | 0.9414 | 0.9484 | 0.9560 | 0.9557 |
| t(LUMO) | 0.7865 | 0.9541 | 0.9460 | 0.9537 | 0.9425 |

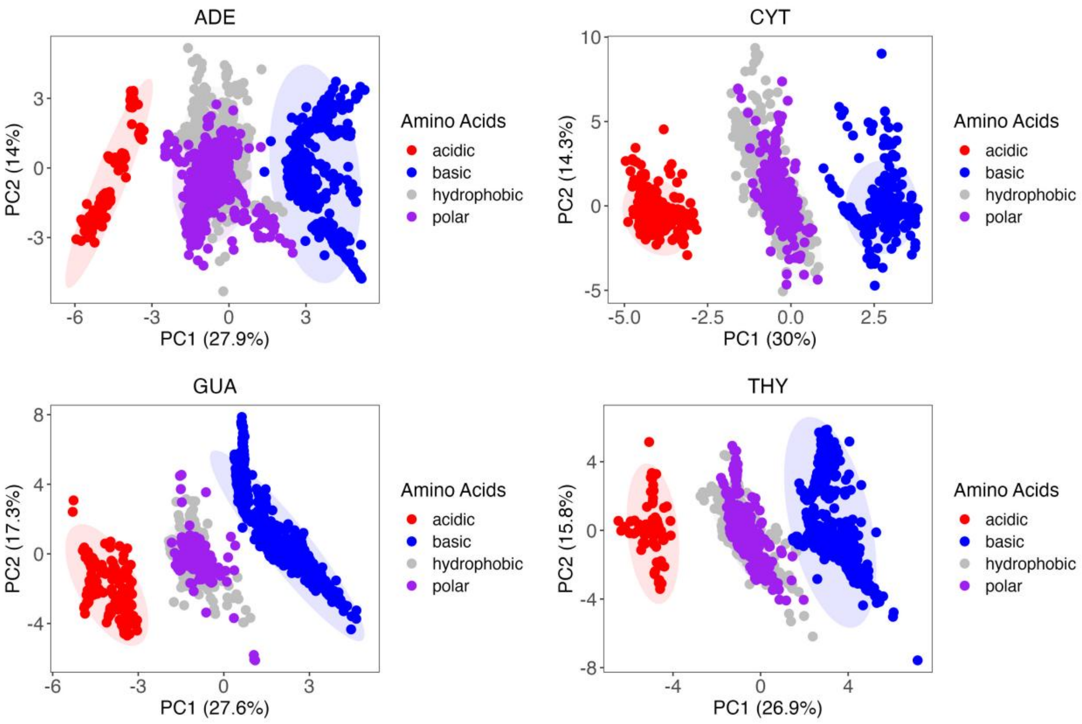


**Figure S1**. The PCA visualization of a spectrum of TB parameters involving HOMO and LUMO orbitals. The TB parameters are generated at B3LYP level.

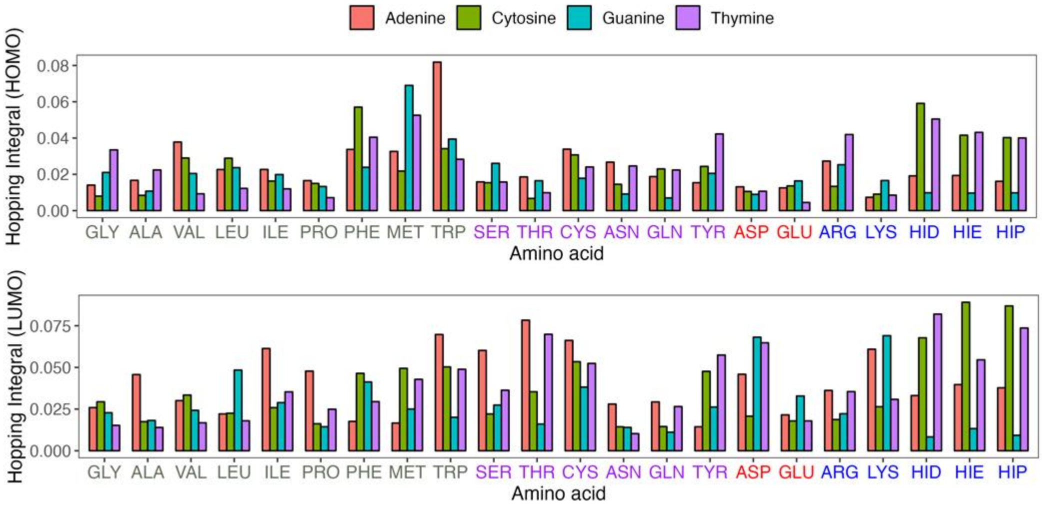


**Figure S2**. The PCA visualization of a spectrum of TB parameters involving HOMO and LUMO orbitals. The visualization includes the four types of nucleic bases, which are the components for any possible DNA sequence. The confidence ellipse represents a statistical probability of 95% that encloses a certain percentage of the data points based on their distribution along the principal components. The TB parameters are generated at B3LYP level.

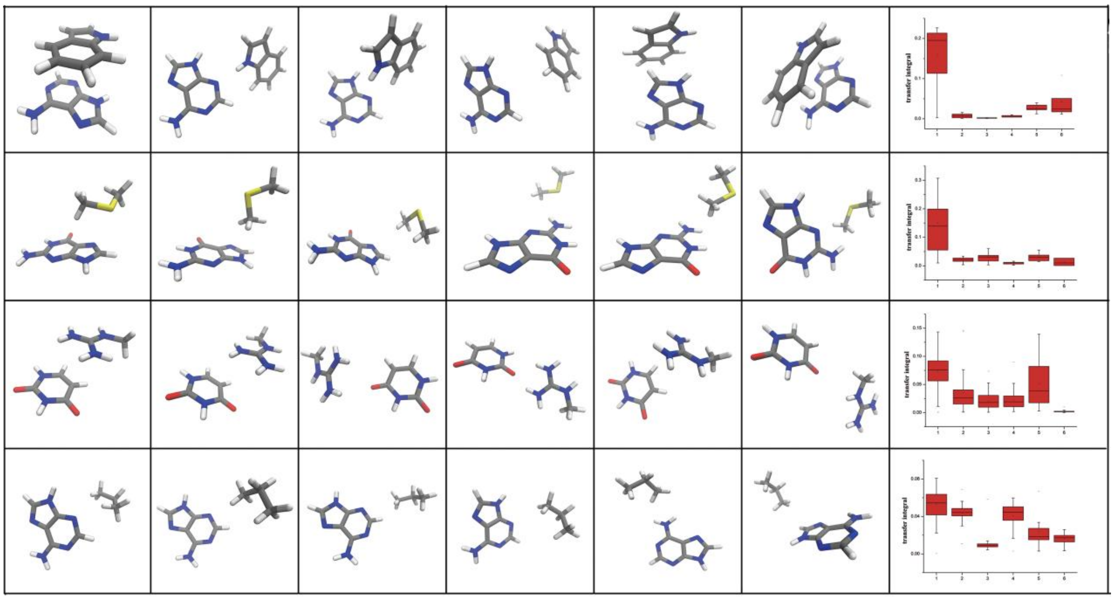


**Figure S3**. The selected structures for four representative sets (ADE/TRP, GUA/MET, THY/ARG, ADE/VAL) of DNA-amino acids pairs with the most significant charge transfer couplings. Spatial distribution of molecules and box plots are used to depict the character of charge transfer couplings.

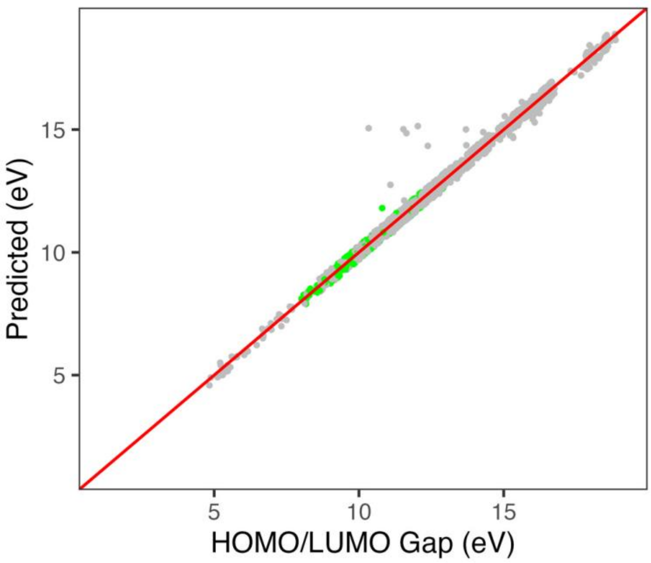


**Figure S4**. Benchmark of the HOMO/LUMO gap for randomly generated dimer and trimer conformers involving nucleobases or amino aicds. The gray dots represent clusters containing only nucleobases or amino acids, while the green dots represent clusters involving both nucleobases and amino acids. The correlation coefficient is 0.9829.

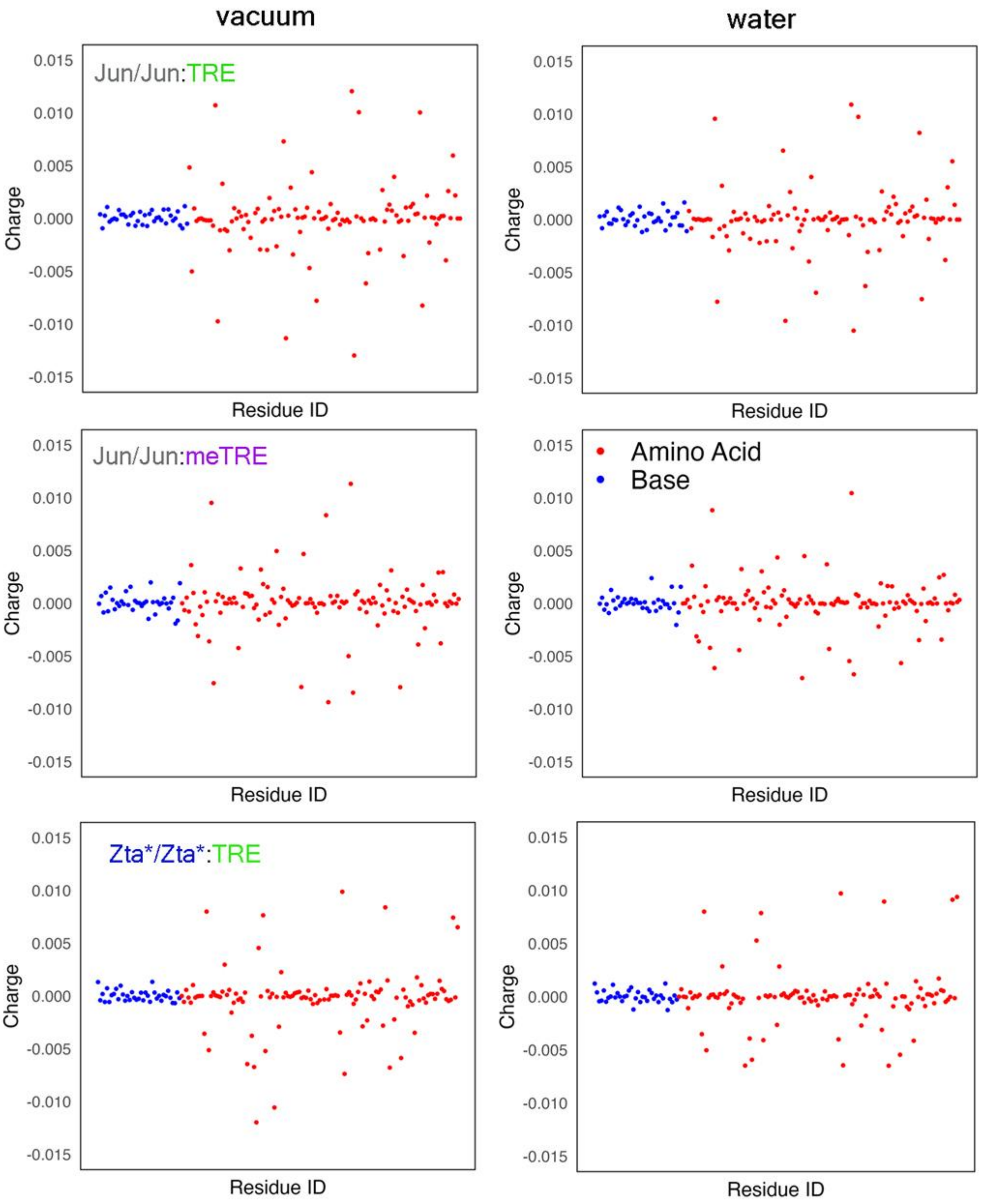


**Figure S5**. The residue level Mulliken charge distribution. Red dots represent the residues from proteins while blue dots correspond to residues from DNA.
